# Finding the strongest gene drive: Simulations reveal unexpected performance differences between *Anopheles* homing suppression drive candidates

**DOI:** 10.1101/2022.03.28.486009

**Authors:** Samuel E. Champer, Isabel K. Kim, Andrew G. Clark, Philipp W. Messer, Jackson Champer

## Abstract

Recent experiments have produced several *Anopheles gambiae* homing gene drives that disrupt female fertility genes, thereby eventually inducing population collapse. Such drives may be highly effective tools to combat malaria. One such homing drive, based on the *zpg* promoter driving CRISPR/Cas9, was able to eliminate a cage population of mosquitoes. A second version, purportedly improved upon the first by incorporating an X-shredder element (which biases inheritance towards male offspring), was similarly successful. Here, we re-analyze the data of each of these gene drives and suggest an alternative interpretation of their performance. We assess each suppression drive within an individual-based simulation framework that models mosquito population dynamics in continuous space. We find that the combined homing/X-shredder drive is actually less effective at population suppression within the context of our mosquito population model. In particular, the combined drive often fails to completely suppress the population, instead resulting in an unstable equilibrium between drive and wild-type alleles. By contrast, otherwise similar drives based on the *nos* promoter may prove to be more promising candidates for future development due to potentially superior performance.

## Introduction

Malaria reduction strategies based on gene drives^1–4^ in *Anopheles* mosquitoes have made substantial advances^5–8^, with several population suppression drives targeting female fertility genes recently proving successful in laboratory settings^5,6^. This raises the possibility that such drives may soon be considered for field deployment. There is thus considerable incentive to engineer the optimal drive for a maximally successful test, which could potentially lead to a more wide-scale field deployment. Because malaria kills over 400,000 people every year while infecting over 200 million^9^, even a modest increase in the efficiency of a gene drive could correspond to a substantial decrease in new cases per year.

Unfortunately, we still lack a complete understanding of the effects of various drive and population characteristics on the outcome of a drive release into a natural population. It is thus difficult to predict which specific type of drive would be most successful under natural conditions. Initial modeling studies assuming a panmictic population indicated that if a homing suppression drive targeting a female fertility gene can avoid the development of resistance alleles that preserve the function of the gene, it can eliminate or at least reduce the population. The exact level of suppression is a function of both species-specific ecological factors and the suppressive power of the drive. This power is often characterized in terms of the drive’s “genetic load”, which for *Anopheles* female-fertility homing drives is typically defined as the fractional reduction in average fecundity of a population in which the drive has reached its equilibrium frequency as compared to an otherwise identical wild-type population. In general, if the genetic load is high enough, panmictic models predict complete population elimination, and any increase in genetic load beyond that threshold can at most provide a decrease in the time to elimination^10^. This suggests that the differences between existing drive candidates that meet this threshold should be minimal.

However, spatially explicit models have indicated that the outcomes of a suppression drive release can be substantially more complicated than those predicted by panmictic population models. In particular, it has been shown that population structure can substantially delay or even prevent complete population suppression for drives with a genetic load high enough to reliably induce population collapse in a panmictic population^11–15^. One mechanism that can prevent population collapse is “chasing”, a phenomenon where wild-type individuals recolonize regions where the drive has eliminated the population. In this situation, they can benefit from the reduced competition due to the low population density in these areas and can substantially increase in number before the drive once again arrives in the area and causes local suppression to reoccur. In this manner, drive and wild-type alleles can persist indefinitely, following a chaotic pattern of local suppression and recolonization. This chasing phenomenon seems to be a common feature of spatial models for many types of suppression drive, regardless of whether the model is implemented using abstract spatial patches^14^, networks of linked demes of mosquito populations^12,13^, or continuous space with discrete generations^11,15^. Unlike panmictic models, these spatial models predict that even modest differences in efficiency between drives can potentially have large effects on the outcome of a drive release, meriting careful consideration of drive candidates to identify those with the greatest potential for success in realistic conditions.

Thus far, several candidates for female-fertility homing suppression drives have been tested in *Anopheles gambiae*. Early drives with the *vasa* promoter offered high germline drive conversion efficiency, but they were not viable candidates due to high levels of somatic Cas9 activity and high rates of embryo resistance allele formation from maternally deposited Cas9 and gRNAs^16,17^. Use of the *zpg* and *nos* promoters was shown to greatly reduce both embryo resistance allele formation and female fitness costs from somatic activity^18^. A drive that combined the *zpg* promoter for Cas9 with a highly conserved gRNA target site in the *dsx* gene (thus preventing the formation of functional resistance alleles) was able to successfully suppress a cage population of mosquitoes^6^. A follow-up study^5^ included a previously developed X-shredder in this drive^19^ to create a male-biased population and was similarly successful in a cage study. Overall, these studies brought forward multiple potential drive candidates that have either already succeeded at suppressing cage populations or which could be expected to do so were they to be implemented with *dsx* as a target.

Here, we assess which of these drive candidates may be most successful at suppressing natural populations of *A. gambiae*. We first obtain multiple possible parameterizations of these drives based on different interpretations of available experimental data. We then analyze them in the context of our previously established individual-based simulation model with continuous space^11^, as well as a new mosquito-specific model with weekly time steps.

## Methods

### Gene drive mechanisms

All of the gene drives modeled in this study are designed to target a female fertility gene that is essential but haplosufficient. Cas9 is directed by a guide RNA (gRNA) to cleave a specific target site located within this gene. Through homology-directed repair, the drive allele then can copy and paste itself into the fertility gene in a manner that effectively inactivates the gene. Since this target gene is haplosufficient, female drive heterozygotes are potentially fully fertile. However, once the drive allele has spread to high frequency in the population, an accumulation of sterile drive-homozygous females will cause the population to be reduced or to completely collapse.

If the cleaved target site does not undergo homology-directed repair but instead is repaired by the error-prone process of end-joining, the resulting mutations may render the site unrecognizable to future Cas9 cleavage. Most of the time, such resistance alleles do not preserve the function of the target gene (these are known as “r2 alleles”) and thus generally do little more than slow the spread of the gene drive. A more severe problem is posed by “r1” resistance alleles, which preserve the function of the target gene and can thereby prevent the suppressive effects of the drive. These r1 alleles were not observed in a cage study of a drive that targeted a highly conserved site^6^, and the use of multiplexed gRNAs could also limit the formation of such alleles^20^. We assume that one or both of these methods will be effective and therefore only model “r2” resistance alleles in our simulations.

One of our drives is motivated by a recent study in which two drives were combined at a single genomic site — a female fertility homing drive and an X-shredder^5^. The X-shredder is designed to cleave multiple sites on the X-chromosome in gametes, rendering them nonviable. This biases the sex ratio towards males because most viable gametes will carry a Y-chromosome. Female gametes that escape destruction may still be sterile since the homing drive disrupts a female fertility gene target as before.

### Discrete-generation simulation model

This model is based on an earlier study^11^ and simulates a population of 50,000 sexually reproducing diploids with non-overlapping, discrete generations using the forward genetic simulation software SLiM (version 3.7)^21^. The wild-type population is allowed to equilibrate for 10 generations before gene drive heterozygotes are released at a frequency such that they represent 1% of the population.

Gene drive processes occur independently in each gametocyte of each reproducing individual. A wild-type allele in a drive heterozygote is converted into a resistance allele with a probability equal to the germline resistance allele formation rate, which differs by sex of the individual. In this study, resistance alleles are always assumed to be “r2” alleles that disrupt the function of the target gene. If the allele remains wild-type, it is converted to a drive allele at a rate equal to the drive conversion rate, which also differs by sex. We also account for maternal Cas9 deposition in the embryo by converting a wild-type allele inherited from a drive-heterozygous parent to a resistance allele with a probability equal to the maternal embryo resistance allele formation rate. In the zpg and zpgX drives (but not the *z*pg2 and zpg2X drives), we account for paternal Cas9 deposition but we assume the effects are mosaic. In this situation, germline drive conversion still occurs assuming the individual is drive heterozygous, but females that suffer from such paternal deposition are rendered sterile.

In this study, we considered drives based on the *nos* or *zpg* promoters for Cas9. The drive could also have an X-shredder (zpgX and zpg2X). However, due to diverging interpretations of existing experimental data, we modeled four possible variations of *nos* drives and two possible interpretations of *zpg* drives. We term these variations nos (without italics), nosF, nosF2, nosF3 (with the latter versions having a lower somatic fitness costs) and zpg (without italics), zpg2, zpgX, and zpg2X (with zpg2 and zpg2X interpreted as not having paternal deposition of Cas9, yet having a higher somatic fitness cost in females). See the results and Table 1 for explanations and parameterizations for each of these drives.

**Table 1.** Drive characteristics.

|  |  |
| --- | --- |
| ***zpg* promoter**<br>Shared characteristics:<br>HDR cut rate in females = 0.99<br>HDR cut rate in males = 0.96<br>Female germline resistance = 0.01<br>Male germline resistance = 0.02<br>Maternal embryo resistance = 0.08 | ***nos* promoter**<br>Shared characteristics:<br>HDR cut rate in females = 0.99<br>HDR cut rate in males = 0.98<br>Female germline resistance = 0.01<br>Male germline resistance = 0.01<br>Maternal embryo resistance = 0.14 |
| ***zpg* drive**<br>Paternal Cas9 deposition rate = 0.69<br>Somatic fitness cost in females = 0.3 | ***nos* drive**<br>Somatic fitness cost in females = 0.45<br>Somatic fitness cost in males = 0.45 |
| ***zpg2* drive**<br>Somatic fitness cost in females = 0.5 | ***nosF* drive**<br>Somatic fitness cost in females = 0.45 |
| ***zpgX* drive**<br>Paternal CAS deposition rate = 0.69<br>Somatic fitness cost in females = 0.3<br>X-shredding rate = 0.93 | ***nosF2* drive**<br>Somatic fitness cost in females = 0.15 |
| ***zpg2X* drive**<br>Somatic fitness cost in females = 0.5<br>X-shredding rate = 0.93 | ***nosF3* drive**<br>No somatic fitness costs. |

### Panmictic model

In the panmictic discrete-generation model, each fertile female randomly selects a mate. We then evaluate the fecundity of the female, which can be reduced by the fitness cost of somatic Cas9 expression. Fecundity is also reduced if the male mate is a nos drive heterozygote (though not in the nosF, nosF2, and nosF3 drives).

Female fecundity (*w*_i_) is further scaled according to a Beverton–Holt model by how close the population size (*N*) is to the carrying capacity of the system (*K*) according to 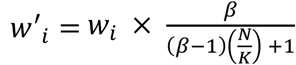, where *β* is equal to the low-density growth rate of the population. We determine the number of eggs produced by drawing from the Binomial distribution, with 50 trials and a success probability equal to 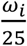 such that a wild-type female is expected to produce 2 offspring on average when the population is at capacity. If the gene drive includes an X-shredder and the father has a drive allele, then the probability that a given offspring will be male is equal to 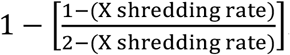. Thus, if Cas9 shreds the X-chromosome of male gametes 100% of the time, all of a male’s offspring will be male, and if the X shredding rate is 0, then the probability an offspring is male is the usual 50%.

Simulations were run until the drive was lost, the population was eliminated, or 1000 generations had elapsed if neither event occurred. In some simulations, modifications were made to facilitate accurate measurement of genetic load (see supplemental methods).

### Spatial model

We extend our panmictic model into continuous space by explicitly tracking every individual’s position across a 1 × 1 (unitless) arena, similar to a model we introduced in a previous study^7^. The simulation begins with 50,000 wild-type individuals that are randomly distributed across the landscape. After 10 generations, a number of drive-heterozygous individuals representing 1% of the total population are released from a 0.01-radius circle at the center of the arena. In the reproduction stage, fertile females can only sample potential mates from a circle surrounding the female with a radius equal to the migration value parameter (with a default of 0.04). If there are no males in this circle, then the female does not reproduce. Reproducing females have a fecundity of 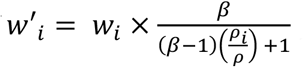, where *ρ*_*i*_ is the local density in a 0.01-radius surrounding the female and *ρ* is the density value that would be expected if the population were evenly distributed across the landscape. This means that a female in a low-density area will have greater fecundity, reflecting potentially greater access to resources as well as reduced competition faced by her offspring. Each offspring is displaced a random distance from its mother. Displacement distance in the *x* and *y* axis are both drawn from a normal distribution with a mean of 0 and standard deviation equal to the migration value parameter. This produces an average displacement of 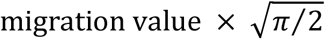. If an offspring’s coordinates fall outside the bounds of the simulation, the coordinates are redrawn until the offspring is placed within the boundaries.

During each simulation, we calculated Green’s coefficient, which provides a quantification of the degree of spatial clustering. To do so, we divided the 1 × 1 area into an 8 × 8 grid and counted the number of individuals present in each of the 64 grid sections. Green’s coefficient (*G*) is defined by 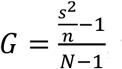, where *N* is the total population size, *n* is mean number of individuals in a grid section, and *s*^2^ is the variance of the counts. If individuals are distributed randomly and according to a Poisson distribution, then it is expected that *n* = *s*^2^ and *G* = 0. By contrast, if all individuals are maximally clustered into a single section of the grid, *G* = 1. Note that we only count wild-type homozygotes in this measurement, because this was found to provide a more useful representation of the spatial dynamics^7^.

As observed previously^11^, the release of a suppression drive can result in a chaotic chase between the drive and wild type alleles. When a suppression drive is first released from the center of the landscape, the population is suppressed radially outward, such that surviving wild-type individuals cluster near the boundaries. This causes Green’s coefficient to increase. However, wild-type individuals that escape into areas previously cleared by the drive are able to produce more offspring as a result of the lack of competition in these areas. If a wild-type population grows into a previously empty area, then Green’s coefficient once again decreases as these individuals occupy more territory. We aimed to capture this inflection point by finding the first local maximum in Green’s coefficient and the first local minimum in the number of wild-type alleles. These events tend to occur within 5 generations of one another. To define the generation when chasing begins in a simulation, we chose the earlier of these two generations.

### *Anopheles*-specific model

In addition to the discrete-generation models, we implemented refined versions of the models that more explicitly simulate the life-cycle, demography, and ecology of *Anopheles* mosquitos. These models progress by weekly time-steps, allowing for overlapping generations.

Mosquito larvae often face fierce competition for resources in the small bodies of water in which they develop while adults do not directly compete with one another for food^22^. Thus, population density no longer regulates adult fecundity in our *Anopheles* model; instead, it affects juvenile survival. In our model, individuals are considered to be in juvenile stages (egg, larvae, and pupae) during their first two weeks of life and reach adulthood when they enter their third week. The larger week-old larvae would not compete with new eggs, and their contribution to competition would stop once they reach the pupal stage. However, they consume more resources compared to smaller larvae and are thus estimated to exert competition at five-fold strength compared to new juveniles. In the panmictic version of this model, newly generated individuals survive until adulthood with a probability that depends on the global sum of new juveniles *n*, the global sum of week-old larvae *o*, the low-density growth rate *β*, as well as the expected competition within the system, which is in turn a function of the number of adult females in a population at capacity (25000 in all simulations) as well as the expected number of offspring per adult female per week (25), calculated as follows:

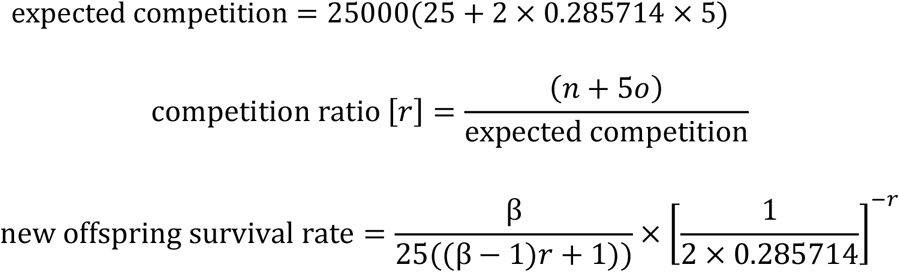

Here, 0.285714 adjusts for the expected number of older juveniles when the population is at equilibrium. Note that while juvenile mosquitoes experience continuous mortality, our model approximates this by determining total juvenile mortality immediately after new juveniles are all generated, representing mortality in both one-week-old and older juveniles (all juveniles that survive this stage will survive to adulthood). This allows more individuals to be culled immediately, thus reducing the computational burden of evaluating the large number of spatial interactions determining competition, thereby allowing for the simulation of larger populations. This approximation is supported by the fact that larvae in their second week are much larger than newly hatched larvae, thus preventing younger larvae from substantially reducing the resources available to older larvae. Later in the second week, the juveniles are pupae, which do not require additional resources. Thus, new juveniles are not likely to affect the mortality of juveniles that are at least a week older, supporting our approximation of determining mortality only in the first week.

In the spatial version of this model, the survival rate of new offspring is affected by the local density of other larvae, rather than global counts thereof. The amount of competition experienced is determined by other new offspring and week-old larvae nearby. The maximum amount of competition contributed by other new offspring is 1.0, which linearly declines to 0.0 at a distance of 0.01 (the average competition contributed by another individual within range is therefore 1/3). Week-old larvae contribute five times as much competition. The expected competition again corresponds to the number of females when the population is at capacity, as well as the expected number of offspring per female:

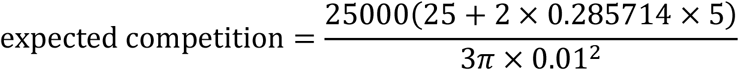

After determining the amount of competition being faced by a new offspring as compared to the expected competition, the survival rate of that new offspring is set in the same way as in the panmictic version of the model.

In the spatial version of the model, surviving adults migrate in the same manner that new offspring are dispersed from their mothers, which is implemented identically to the discrete spatial model except that the migration value now represents the average displacement per week, with the default of 0.0307 creating an equal displacement per generation (males and females together averaging 2.67 displacements in a generation) as well as an equal “drive wave advance speed” to the discrete generation model default of 0.04 (with an average displacement of 0.05).

Comparing our default dispersal rate of 0.307 to a recent study that found a mean dispersal of 171 m over 20 days^23^, we can potentially state that our mean dispersal corresponds to 101 m per week and that our simulation area length is therefore 3.3 km. A population density of 430 mosquitoes per hectare would thus yield a total population of 470,000 adults, compared to our default of 40,000 (of which 25,000 are females). However, the effective population (more akin to what is generally analyzed in population genetic models) would be substantially below 470,000, and there is certainly substantial density variation between wild mosquito populations. Thus, our modeled population density could be considered reasonable when modeling *Anopheles*, based on length scales from our migration rate. However, it is unclear what parameter value for the density interaction radius would best represent actual mosquito populations, given the varying size and distribution of larval habitat, as well as larval movement. Thus, our value of 0.01 stands in as an estimate, chosen as a low value that is still large enough to avoid extreme variation in individual larval competition when the population is at equilibrium before a drive release.

Female mosquitoes of most species usually just mate once, though older females have often been observed to mate a second time^24–27^. Females that have already mated store sperm from the male, which is used to continue to fertilize eggs in future weeks. This behavior is implemented in the model. We also implemented a 5% chance each week that the female will re-mate, resulting in the new mate fathering any future offspring unless re-mating occurs again. This 5% weekly re-mating value is based on estimates from *A. gambiae* experimental^27^ and field^26^ studies, assuming lower field survival rates.

After reaching adulthood, females have a 50% likelihood to successfully produce offspring during any given week if they have previously mated (which takes place before offspring production). The number of eggs laid is not density dependent but is instead drawn from a Poisson distribution with the average set at 50 times the product of the fitness of the two parents. The number of eggs laid by *A. gambiae* appears closer to three times this level in laboratory conditions^28^, but in practice, usually only a far smaller number reach adulthood in wild conditions. Our use of 50 allows larva at low density to have extremely high survival rates compared to most wild conditions while still minimizing computational burden.

The survival rates of adults are also not density dependent. *Anopheles* females have longer lifespans than males. In our model, males never survive beyond their fifth week and females never beyond their eighth, with the survival rates at each age of adulthood given as follows:

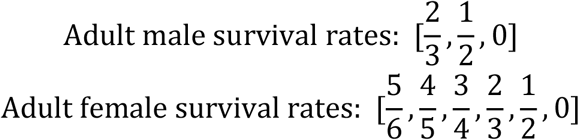

This results in an approximately linearly declining number of surviving adult members of a single week cohort. The shape of the curve was chosen based on survival curves in laboratory studies^29^ and to allow simulation of age-based health. The measured male survival rate in the field was measured as approximately 30% per week in one recent study^23^, and often higher in females and closer to 50% in males for older studies^30,31^. This is somewhat lower, but broadly similar to our model, considering high variation in field conditions and our need to simulate a high effective population size with limited computational resources.

Note that individuals have the opportunity to mate and reproduce before these survival rates take effect. One generation in the discrete model is then equivalent to ~3.167 weeks in this model (thus, we ran these simulations for 3,167 weeks to represent 1,000 generations).

### Data generation

Simulations were run on the computing cluster at the Department of Computational Biology at Cornell University. Data processing and analytics were performed in Python, and figures were prepared in Python and R. All SLiM files for the implementation of these suppression drives are available on GitHub (https://github.com/jchamper/ChamperLab/tree/main/Mosquito-Drive-Modeling).

## Results

### Parameterization of *Anopheles* suppression gene drives

Several suppression drives have been constructed by the Crisanti lab in *Anopheles gambiae*^5,6,17,18^. By examining these data, we calculated drive conversion rates (the rate at which drive alleles are converted to wild-type alleles in the germline) in female and male drive/wild-type heterozygotes, and we similarly obtained estimates of the germline resistance allele formation rates (distinctly in females and males), embryo resistance allele rates from parental deposition of Cas9 and gRNAs (both maternal and paternal), and additional fitness costs (see Supplemental Results for details). We assumed these fitness costs were from somatic expression of Cas9 and gRNA in drive/wild-type heterozygotes, with no intrinsic fitness cost of the drive itself (such costs appear to be small based on previous studies^7,8,32–34^). Table 1 contains parameterization of each modeled drive.

The first drive to demonstrate suppression of a cage population targeted a conserved site of *dsx* using Cas9 expressed by the *zpg* promoter^6^, which was also assessed in a drive with another target site^18^. We model the authors’ interpretation of this study with paternal Cas9 deposition (zpg) and another interpretation that instead assumes no paternal deposition and higher somatic fitness costs (zpg2), which we consider more likely (Supplemental Results).

A follow-up study by the same group added the I-PpoI nuclease to the drive, thus allowing it to shred the X-chromosome and bias the population toward males^5^. According to their data, 93% of X-chromosomes are effectively shredded in the germline. We model this variant of the drive with both the original and alternate parameter sets for the *zpg* suppression drive (zpgX and zpg2X, respectively).

The *nos* promoter has also been shown to support highly efficient homing suppression drives. We parameterize this drive (nos) based on a previous study^18^ with three additional alternative interpretations of the data (Supplemental Results). In one (nosF), we assume no effect of somatic Cas9 expression in males, which we believe may be more realistic. A second possible alternative interpretation assumes reduced somatic effects in females (nosF2), and a third highly optimistic interpretation assumes no somatic effects (nosF3).

### Drive performance in the panmictic discrete-generation model

We first simulated the drives in our panmictic discrete-generation model to assess their basic properties, starting with genetic load. Genetic load describes the reduction in reproductive capacity of the population compared to a population that is identical except for being composed entirely of wild-type individuals. In panmictic populations, this measurement often reaches an early peak as the drive allele reaches its maximum frequency, but then slightly declines to a steady value once the drive allele and nonfunctional resistance alleles reach their equilibrium frequency. The rate at which wild-type alleles are converted to drive alleles in the germline of drive heterozygotes is the main determinant of genetic load. Negative fitness effects associated with the drive can reduce the genetic load, as can the rate at which nonfunctional resistance alleles are formed in both the germline and embryo, though the effect of such alleles is usually not large^35^. To eliminate a panmictic population, the drive must induce a sufficiently high genetic load in order to overpower the growth of wild-type populations at low density. All of the implementations of the *nos* drive were found to have a higher genetic load than the *zpg* drives (Figure 1), though zpg2 performed well compared to the other *zpg* drives. Notably, the addition of an X-shredder was detrimental to zpg2, with zpg2X’s genetic load reduced by 0.16. Even the highest fitness cost interpretation of *nos* showed a genetic load of 0.96, with the lower fitness cost interpretations scoring even higher. Three of the drives with low equilibrium genetic loads (zpg, zpgX, and zpg2X) actually had higher peak genetic loads shortly after their release, with the genetic load eventually declining due to formation of nonfunctional resistance alleles.

**Figure 1.**
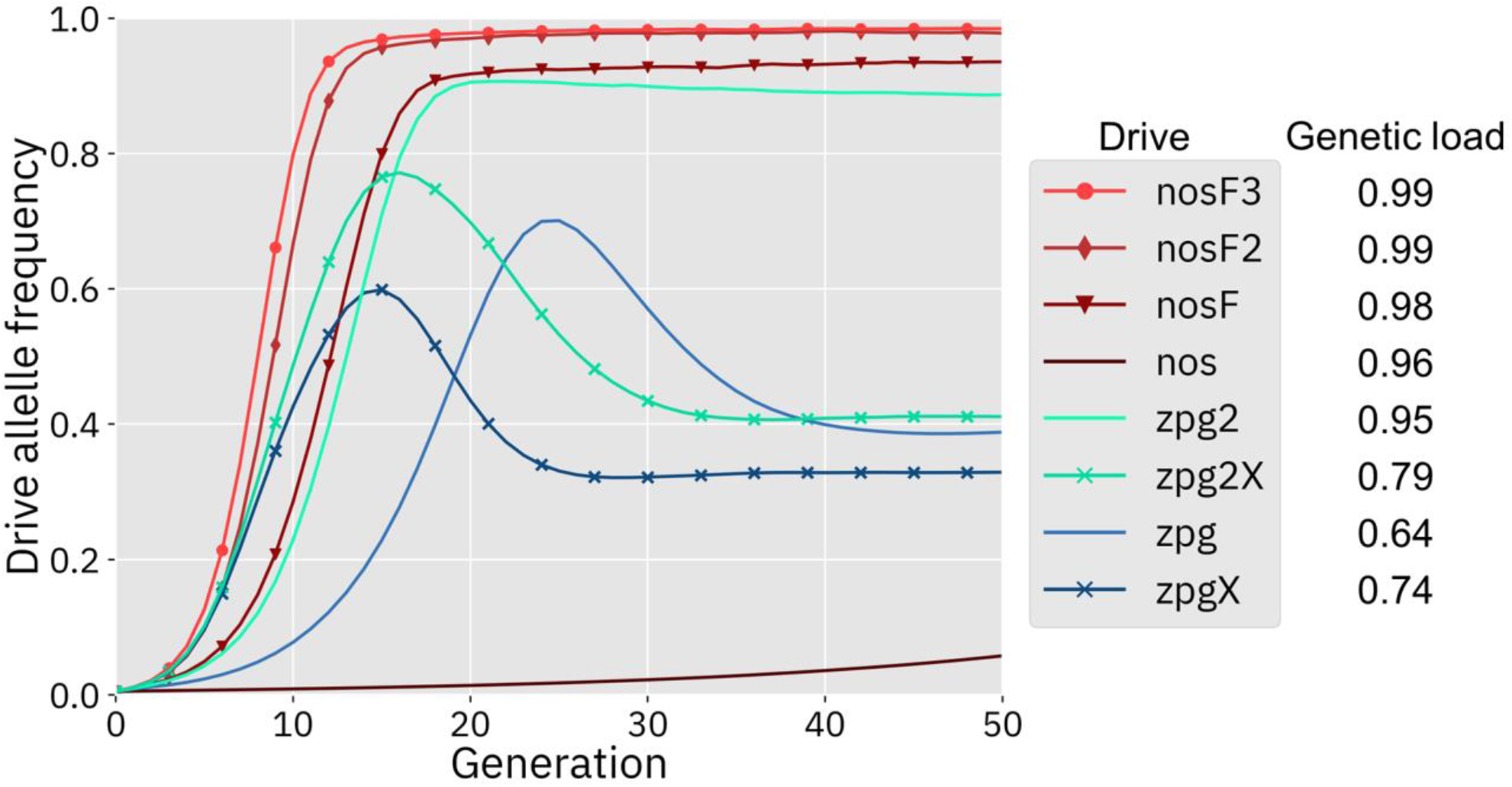
Drive allele frequency trajectories in the panmictic discrete-generation model. Using default parameters, each drive was released into a panmictic population at 0.5% initial frequency (1% heterozygote release). The average allele frequency as estimated over 100 replicates per drive is displayed for each generation. Offspring were artificially generated from fertile individuals at high rates to prevent complete population suppression even at high drive frequencies and genetic loads. For a description of this method of measuring genetic load, see the Supplemental Methods.

Next, we measured the rate at which the drive spread through the population (Figure 1). The addition of the X-shredder substantially improved the speed of both *zpg* drives, with the zpg2 interpretation resulting in considerably higher performance than zpg. The nos drive, with its high fitness cost in both males and females, performed far worse than all of the other drives in this regard, seemingly belying its high genetic load; the somatic fitness costs had so great an impact on fecundity that the drive could make little headway given a low starting frequency (though the drive still eventually reaches a high equilibrium frequency, see Figure S1). However, the other *nos* drives performed well, particularly nosF2 and nosF3. Even nosF outperformed zpg2, though both *zpg* drives with X-shredders were still faster.

### Spatial discrete-generation model

Previously, we found that drive outcomes in a model with continuous space can substantially differ from those in panmictic populations^11^. In our spatial discrete-generation model, the migration value controls the radius in which a female can find a mate as well as the displacement between a mother and her offspring. The low-density growth rate is a multiplier on female fecundity in the absence of competition. To examine how each drive would behave under various ecological assumptions, we varied these two parameters and recorded whether the simulations resulted in (a) suppression without chasing, (b) suppression after chasing, (c) long-term chasing (defined as 1000 or more generations of chasing), (d) drive loss after chasing, or (e) drive loss without chasing.

As observed in our earlier study on suppression gene drives in continuous space^11^, the low-density growth rate had a large impact on drive performance. Figure 2 and Figure S2 display the result of 200 simulations at realistic parameter values for each drive (with a low-density growth rate of 10 and 6, respectively). Under both parametrizations, the zpg drive was unable to fully eliminate the population; instead, long-term chasing occurred in 100% of simulations. However, our alternative parameterization of the drive, zpg2, achieved suppression 28% of the time when the low-density growth rate was 10 (Figure 2A) and 63% of the time when the low-density growth rate was reduced to 6 (Figure S2A). Similar to zpg, the zpgX drive resulted in long-term chasing in almost all simulations, though the average number of fertile females was substantially reduced. However, our alternative parametrization, zpg2X, was less likely to result in long-term chasing, and suppression was observed in 12% of outcomes when the low-density growth rate was 10 (Figure 2A) and 23% of outcomes when it was reduced to 6 (Figure S2A). This is a substantially lower rate than the zpg2 drive, but the average number of fertile females during chases was also somewhat lower.

**Figure 2.**
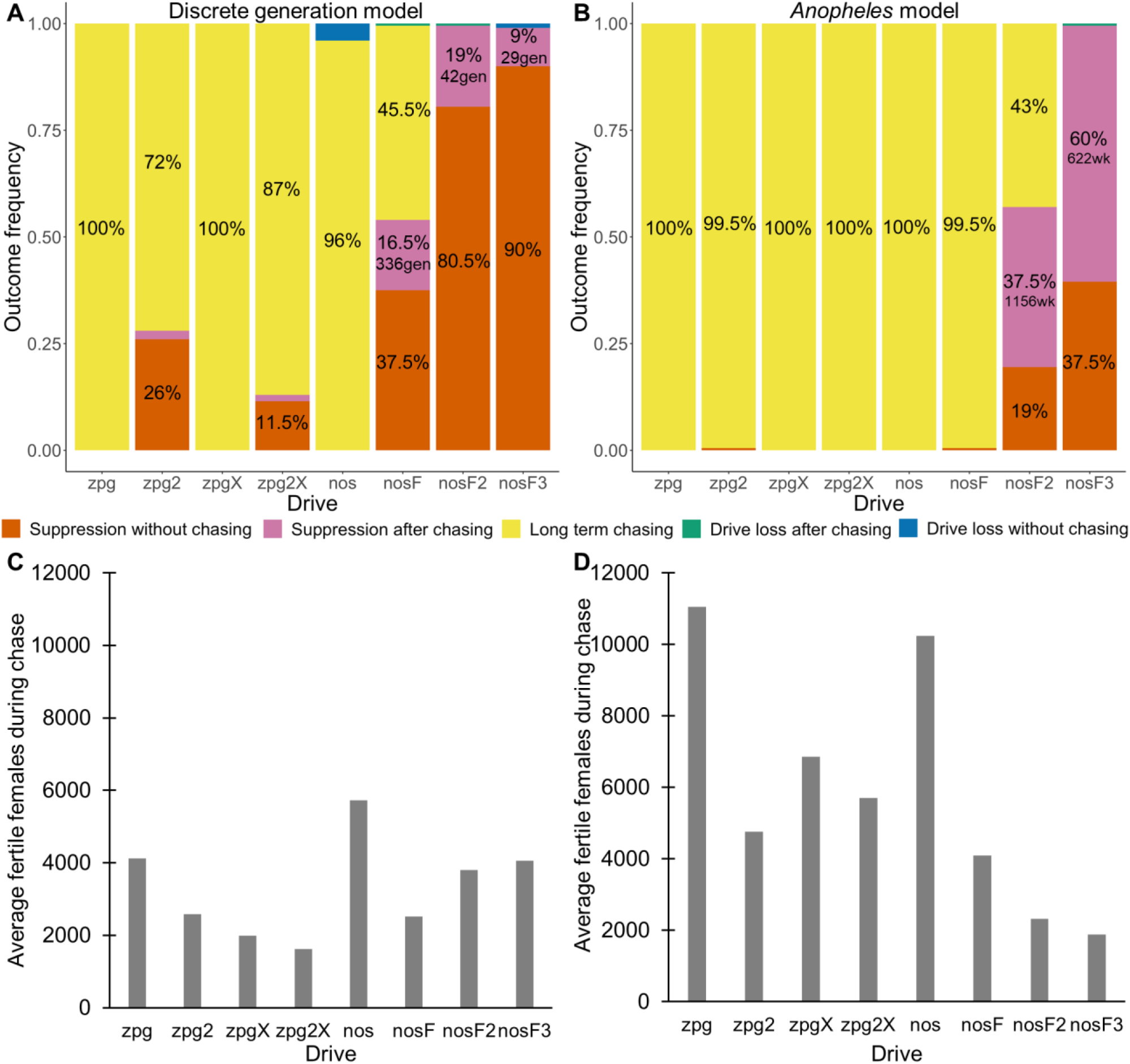
Outcomes in the spatial models. Using default parameters, a low-density growth rate of 10, and with 200 replicates per drive, each drive was released into the middle of a wild-type population. The outcome was recorded after 1000 generations or when the population was eliminated for the discrete-generation (**A**) and *Anopheles*-specific (**B**) models. In outcomes involving chasing followed by suppression, the number of generations (gen) or weeks (wk) between the start of chasing and population elimination is shown. Also displayed is the average number of fertile females during periods of chasing (including both long-term and short-term chasing outcomes) for the discrete-generation (**C**) and *Anopheles*-specific (**D**) models. Due to the high number of replicates, the error for each data point is negligible, except for the nosF2 and nosF3 drives in the discrete-generation model due to the short duration of chasing.

The standard nos drive resulted in long-term chasing in almost 100% of simulations with both low-density growth rate values (Figure 2A, Figure S2A). However, when the somatic fitness cost in heterozygote males was omitted (nosF), drive performance dramatically improved. When the low-density growth rate was 10, nosF achieved suppression in 54% of simulations (Figure 2A), and when it was lowered to 6, the suppression rate increased to 89%, a notable improvement over the best performing *zpg* drive. With a decreased female heterozygote somatic fitness cost in the nosF2 and nosF3 drive, suppression was almost guaranteed.

To further examine the behavior of these drives, we varied the migration rate from 0.01 to 0.06 and the low-density growth rate from 2 to 12. The results were broadly consistent with the results of our earlier study^11^. Figure 3 shows the most common outcome from the simulations, and Figures S3-6 show the likelihood of each outcome. In general, suppression tended to be increasingly likely at higher migration values and when low-density growth rates were smaller (Figure S3). There was usually a “transition” regime (Figure 2A) involving suppression after chasing (Figure S4) in between rapid suppression outcomes and long-term chasing outcomes (Figure S5). Drive loss usually only occurred when migration values and low-density growth rates were both very low (Figure S6). At higher migration values and lower low-density growth rates, “suppression after chasing” became more limited in duration (Figure S7), and the average number of fertile females was reduced even during long term chases (Figure S8), which would likely substantially reduce disease transmission.

**Figure 3.**
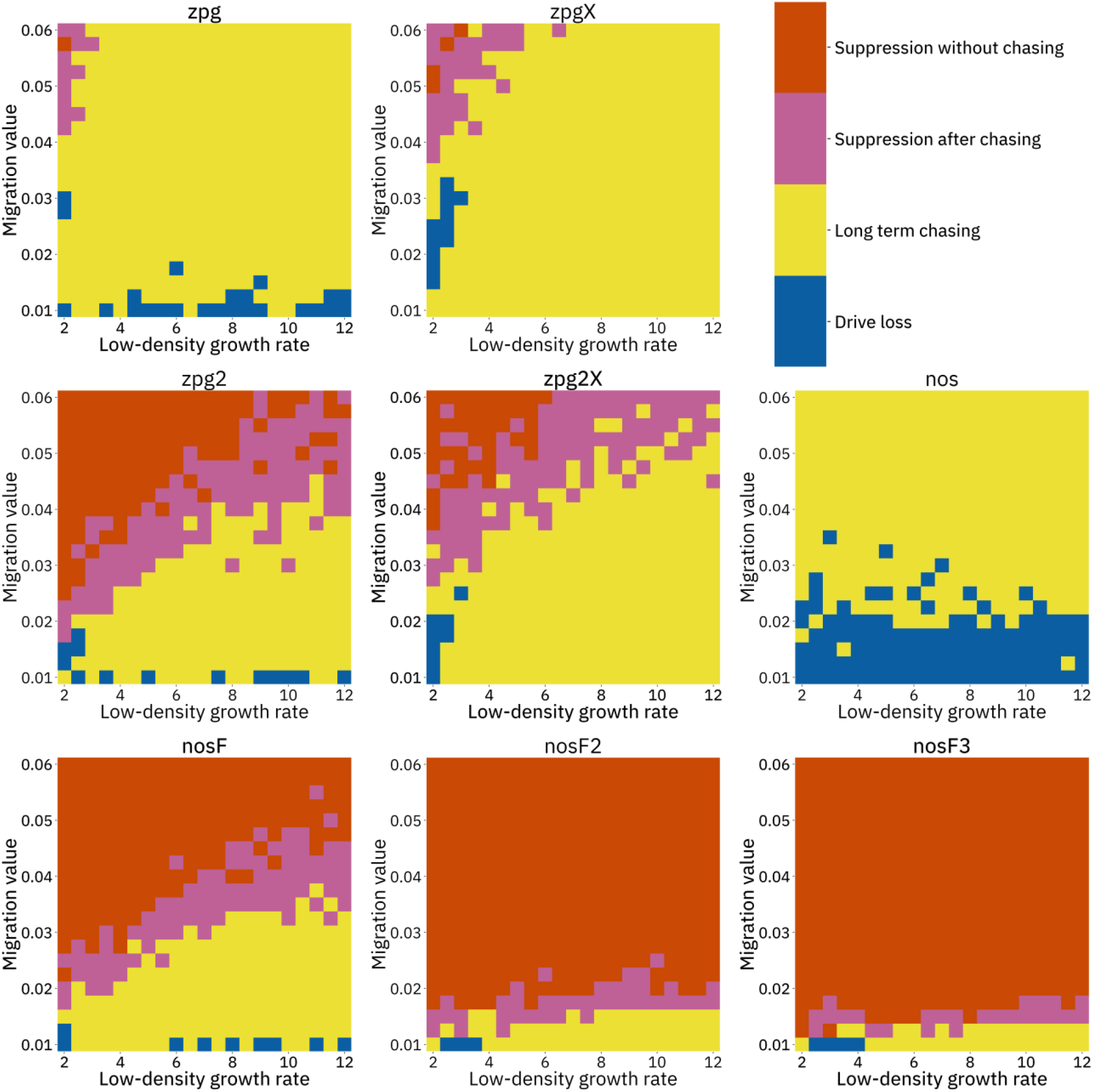
Impact of low-density growth rate and migration on outcomes in the discrete-generation model. The color of each square represents the most common outcome from 20 simulations, after adjustment to show the most representative outcome. The adjustment counts pairs of “suppression without chasing” and “long-term chasing” outcomes as two instances of “suppression after chasing”.

Overall, the three intermediate performance drives (zpg2, zpg2X, and nosF) had outcomes that depended heavily on the migration value and to a lesser extent on the low-density growth rate (Figure 3). The strongest drives, nosF2 and nosF3, were able to induce suppression over most of the parameter space, while release of the three weakest drives, zpg, zpgX, and nos, usually resulted in long-term chasing outcomes.

### *Anopheles*-specific model

In addition to our discrete-generation model, we implemented a model that more explicitly simulates the expected dynamics of an *Anopheles* population by modeling overlapping generations using week-long time steps. The panmictic version of this model was used to calculate the genetic load of each drive as well as the speed at which it was able to spread, in the same manner as the discrete-generation model. The genetic load values measured by both models was within 1% for all of the drives (Figure 1). The allele frequency trajectories varied slightly between the two models, but only for drives with X-shredders that biased the sex ratio (Figure S1). This occurred because the drive was mostly present in males, which have a shorter adult-stage lifespan than females, thus reducing the overall drive-allele frequency even though frequencies in emerging adults was the same.

The drives were next assessed in the spatial version of the *Anopheles*-specific model (Figure 2B). Generally, long-term chasing outcomes were more frequent in these simulations than in the discrete-generation model (Figure 2A). This came with an accompanying decrease in suppression outcomes as well as drive-loss outcomes. For example, in the discrete-generation model with a low-density growth rate of 10, zpg2 was able to suppress the population in 28% of simulations, while the drive suppressed in only 0.5% of simulations in the mosquito model. The average number of fertile females during chasing increased from 2567 to 4760. In the case of nosF, performance was similarly decreased from suppression in 54% of simulations to only 0.5% of simulations when the low-density growth rate was set to 10, with the average number of fertile females during chasing increasing from 2498 to 4101. In both models, nosF2 and nosF3 retained the ability to suppress the population. However, in the *Anopheles*-specific model, suppression often occurred only after an initial chase, and only NosF3 could still reliably suppress the population in almost every simulation (Figure 2B). The zpg2X drive also performed notably more poorly in this model. In the discrete-generation model, the drive was able to suppress the population in 10% of simulations, but in the *Anopheles*-specific model, the drive was only able to suppress 0.5% of the time. The reduction in the number of fertile females achieved by both X-shredder drives in this model was also less pronounced and, notably, was inferior for zpg2X as compared to zpg2. This was perhaps because the reduced genetic load of zpg2X outweighed the advantage of the male-bias. All the drives had similarly increased performance when the low-density growth rate was adjusted to 6 (Figure S2B), This allowed nosF2 to more reliably suppress the population, and the average number of fertile females was lower for all drives.

We next varied the low-density growth rate from 2 to 12 in steps of 1.0, while varying the migration value from 0.008 to 0.046 in steps of 0.0038 (thus corresponding to the same range that was examined in the discrete-generation model). The above tendencies mostly held true in this analysis as well (see Figure 4 and Figures S9-14). Chasing outcomes tended to replace suppression outcomes and dominate large portions of the parameter space (Figures S9-11, compare with Figures S3-5). Drive-loss outcomes were extremely uncommon in this model compared to the discrete-generation model, with such outcomes occurring in significant quantity only when the low-density growth rate was very low (Figure S12, compare with Figures S6). Only nosF2 and nosF3 performed well in this model and were able to suppress the population at moderate to high migration values (Figure S9, compare to Figure S3), or at least keep the duration of chasing and average number of fertile females at low levels (Figure S13-14, compare to Figure S7-8).

**Figure 4.**
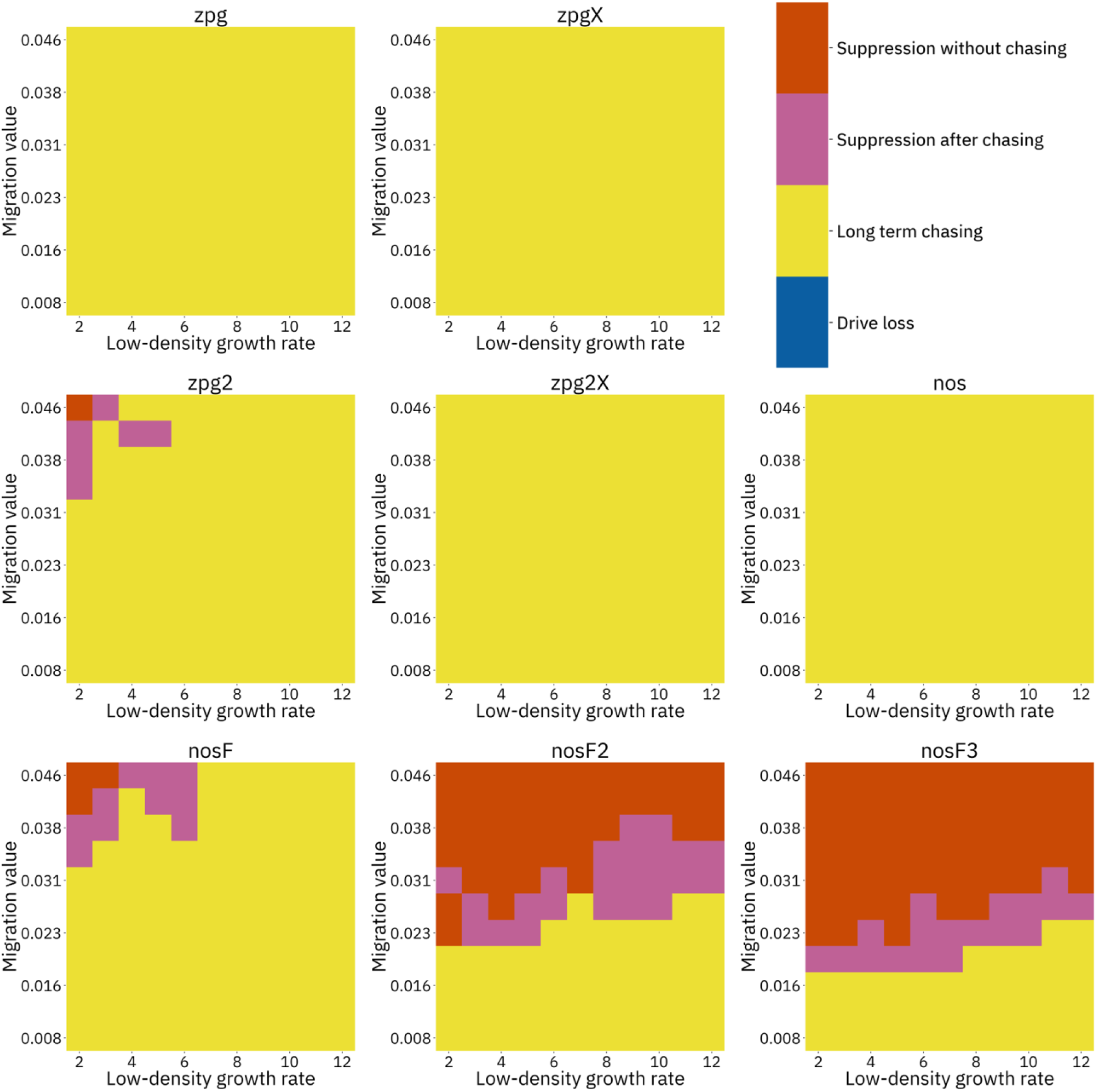
Impact of low-density growth rate and migration on outcomes in the *Anopheles*-specific model. The color of each square represents the most common outcome from 20 simulations after adjustment to show the most representative outcome. The adjustment counts pairs of “suppression without chasing” and “long-term chasing” outcomes as two instances of “suppression after chasing”. Note that the range of migration values in this model correspond to the same net migration per generation as those found in the range of in Figure 3.

## Discussion

In this study, we analyzed several possible types of homing gene drives for population suppression of the malaria vector *Anopheles gambiae*. Various parameter sets for each of these drives were considered, representing alternative explanations for fecundity measurements previously conducted^5,6,18^. We found that the implementation of a drive involving a combined homing drive and X-shredder may actually be less capable of suppressing a population than the original *dsx* drive with the *zpg* promoter alone, particularly in a spatially continuous population. Further, we found that the *nos* promoter may be a preferable alternative for Cas9 expression compared to the *zpg* promoter, depending on which interpretation of the experimental data proves to be the most accurate.

These differences in interpretation of data could perhaps be cleared up with additional experiments. For example, the rate of paternal Cas9 deposition for a given Cas9 promoter could be assessed using a split-drive system^36^, sequencing, or conducting fecundity assessment of offspring that have a drive-carrier father but that do not themselves inherit a Cas9 allele. Though the exact rate may be somewhat different due to differences in expression at distant genomic loci, such an experiment should still serve to confirm the presence and magnitude of this phenomenon. Batch effects could also have large effects on fecundity measurements^18^, which we have also noticed in our *Drosophila* experiments^32,33,37,38^. To address this, fecundity experiments should, when possible, be performed with individuals that were raised in the same container (and then subsequently separated by fluorescence, ideally for multiple generations) and preferably even from the same parent (though this is only a readily available option for split-drive systems, especially when drive conversion efficiency is very high, as achieved by most *Anopheles* drives). Such experiments could potentially reveal that the negative fitness effects of the *nos*-Cas9 drive are smaller than initial results indicated. The *nos* promoter still needs to be assessed at the *dsx* target site, but this may prove to be an excellent combination if drive conversion remains high and resistance rates low.

Based on our modeling results, it appears that a combined X-shredder system may not represent an improvement to the standard homing suppression drive for a couple of reasons. First, with the alternate parameter set, the genetic load of this drive is lower (even with perfect homing, the 93% shredding rate would yield a genetic load of only 0.87) and is perhaps not high enough to reliably suppress a robust wild population. Second, the drives including the X-shredder do not seem to perform well in spatial simulations, even with higher shredding rates^11^. This held true in our *Anopheles*-specific simulations, where the release of these drives usually resulted in long-term chasing with a high average number of fertile females. However, combined X-shredder systems are not without advantages. They can provide a way to support some continued suppression even in the presence of functional resistance alleles (though the rate that such alleles form at *dsx* is unknown and may already be sufficiently low due to the highly conserved target site, which is essential at the sequence level^11^, and it could possibly be further reduced by the use of a drive with multiplexed gRNAs^20^). Also, because X-shredders bias the population toward males, such systems may effectively reduce biting females (and therefore reduce disease transmission) more quickly than standard homing suppression drives^5^.

In general, the results of our mosquito-specific continuous space model mirror the outcomes of our earlier generic model^11^ of suppression gene drives in continuous space, but the drives have substantially greater difficulty eliminating the population. Indeed, in the *Anopheles*-specific model, chasing outcomes were substantially more common, suggesting that chasing may be difficult to avoid when using a gene drive in natural populations unless the drive meets stringent efficiency criteria. These differences between models were likely at least partially due to the high reproductive capacity of *Anopheles* mosquitoes, which ensures a robust population even under high genetic load due to the reduction in larval competition when fewer individuals are reproducing. The effective migration level of the drive allele into wild-type populations is also reduced by way of females only mating once, and such reduced migration also tends to promote chasing^11^. It should be noted that while our *Anopheles*-specific model represents a step toward increased realism compared to a discrete-generation model, it still has various limitations. For example, distances were still unitless (though see methods for possible comparison to previous field studies), and higher relative dispersal rates in real populations may enable easier suppression. We did not consider the possibility of long-distance migration, which could make chasing more likely, or functional resistance, which can evolve more readily during lengthy chases^11^. We also did not take into account other factors that may be important in real-world populations, such as seasonality, variation in survival rates due to climate or predation, heterogeneous landscapes, nonrandom movement, genetic variation, or the possibility of evolutionary responses to the spread of a suppression drive in the population^11,14,39,40^.

One potentially important implication of these models is that many of the gene drive designs currently under consideration may be quite close to the boundary between drives capable of successful suppression and drives expected to fail due to chasing, such that the outcome of a drive release could be very sensitive to the precise ecological characteristics of the targeted population. Small parameter differences in drive performance could therefore be critical in ensuring drive success, suggesting that currently ambiguous drive characteristics should be reconsidered and measured with as much accuracy as possible. At the same time, computational models must be further refined in order to improve their predication accuracy, so that we can more reliably assess whether a candidate suppression drive indeed meets any specific requirements for success, or to at least understand the most likely outcome.

In conclusion, our modeling indicates that several promising candidates for suppression gene drives could have substantially different overall effectiveness in spatial mosquito populations than predicted by panmictic population models. Specifically, we believe that a standard homing suppression drive using the *nos* promoter^8,18^ for Cas9 (possibly a high fidelity variant to avoid off-target cleavage^41^) and targeting *dsx*^5,6^ with two or more closely spaced gRNA target sites^20,38^ may be the optimal combination of currently available tools for population suppression of *Anopheles* mosquitoes.

## Acknowledgements

This study was supported by the National Institutes of Health award F32AI138476 to JC and award R01GM127418 to PWM.

## Supplemental Information

### Supplemental Methods

#### Measurement of genetic load in panmictic simulations

A subset of panmictic simulations were used to assess the genetic load of the drive (the reduction in reproductive capacity of the population compared to a population that is identical except for being composed entirely of wild-type individuals). For sterility-based homing suppression drives, this measures the reduction in fecundity caused by the presence of the drive. As measured at any generation *t* in the simulation, the genetic load is a function of the number of females (*N*_*f*_) actually alive in the following time step and the number that could be predicted if there were no drive present:

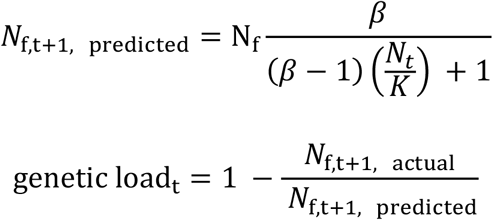

However, genetic load as measured in this manner is merely an instantaneous measurement – it describes the impact a drive has on the population at only a single time step. Further, without modification, it is not possible to simply average the genetic load measured at several generations in a row, because the population can cease to exist under the pressure of the drive before sufficient measurement can be made. To remedy this, in simulations specifically seeking to determine genetic load, fertile females had a multiplier applied to the number of offspring they generated in order to approximately maintain the population at capacity given the number of infertile individuals in the system at any given time. Specifically, the number of offspring produced by fertile females was divided by the following population factor:

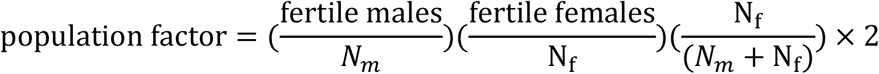

Similarly, the numerator term of the measured genetic load was multiplied by this factor in order to account for the excess individuals beyond the number that would normally be present in the next generation.

## Supplemental Results

### Parameterization of drive candidates

We first examined drive inheritance from *zpg* promoter drive heterozygous individuals from two studies^6,18^. Discounting males that did not show biased inheritance (such males likely had a resistance allele form while they were embryos due to maternally deposited Cas9), the drive conversion rate appears to be 96% for male heterozygotes and 99% for female heterozygotes. We set the germline resistance allele formation rate to be 2% for males and 1% for females so that approximately half of the alleles that fail to undergo drive conversion remain wild-type (these estimates don’t substantially impact our results since we make the assumption that all resistance alleles disrupt the function of the drive). Males with drive mothers (who therefore receive maternally deposited Cas9 and gRNA) that failed to show substantial drive conversion in the germline likely had embryo resistance alleles, as would sterile females with drives mothers, so we posit an embryo resistance allele formation rate of approximately 8% based on these frequencies. The authors hypothesized that reductions in female fertility could also be due to paternal deposition leading to resistance allele formation^18^. Based on the rate of sterile female drive carriers with male drive parents, we calculated the paternal resistance allele formation rate as 69%. However, unlike maternal resistance alleles, we consider these paternal resistance alleles to be mosaic at a sufficient level to cause complete female sterility, but not present at sufficient level in the germline to prevent efficient drive conversion (thus, these resistance alleles have no effect on male progeny, in line with experimental data). We then applied a 30% fitness cost in all drive/wild-type heterozygous females to account for the remainder of the reduced fertility of drive females compared to wild-type females, though this reduction could also be interpreted as additional mosaic parental deposition from both drive mothers and fathers.

However, alternative explanations exist for the reduced fertility in female drive heterozygotes with male drive parents as compared to those with female drive parents. These reduced fertility levels could be caused by batch effects on fecundity or simply be random variations. If interpreted in this manner, these fecundity reductions can be explained by somatic Cas9 expression and cleavage, with no significant paternal deposition. This is supported by another data set with a different target gene but identical drive components, wherein female heterozygotes had similar fecundity regardless of which parent provided the drive allele^18^. Fundamentally, it seems unlikely that paternal deposition causes more embryo resistance allele formation than maternal deposition, considering the relative size of the sperm and the egg. The fact that the zpg drive targeting *dsx* completely suppressed a cage population is also indirect evidence against the drive being so negatively impacted by paternal deposition. We found that this interpretation of the drive has a relatively low genetic load, which would perhaps be insufficient to suppress a cage population (see Table 1). Specifically, a single fully fertile female mosquito appeared to be able to generate 130 viable larval progeny based on individual crosses^6^. This could mean that 650 eggs (the number used for each generation of the cage) could potentially be generated by as few as five fully fertile females. Even conservatively assuming that 650 eggs yields only 150 adult females, a genetic load of at least 0.97 in a deterministic model is needed to reduce the viable egg count of the next generation to below 650 on average. Though stochastic effects could certainly result in a cage population being suppressed by a drive with a somewhat lower genetic load, this explanation is increasingly unlikely at lower genetic loads. Finally, previous studies have found no indication of paternal Cas9 deposition in *Drosophila*^20,36,42,43^ and nothing conclusive in *Anopheles*. Evidence of the phenomenon found in *Anopheles* studies^17,44^ could also potentially be similarly explained as effects caused by leaky somatic Cas9 expression (though significant paternal deposition has been clearly observed with a different nuclease^19,45^). In our models, zpg2 is parameterized to match this alternative explanation. Instead of a 69% paternal embryo resistance allele formation rate, this implementation has a 50% fitness cost from somatic cleavage in female heterozygotes (increased from 30%, thus matching the average fecundity reported in two studies^6,18^).

The *nos* promoter represents a potential alternative to *zpg*’s germline-restricted expression of Cas9. We parameterize this drive based on a previous study^18^ using 99% drive conversion in female drive/wild-type heterozygotes, 98% drive conversion in males, 1% germline resistance allele formation for both sexes, and a 14% embryo resistance allele formation rate in the progeny of female drive heterozygotes. In the previous study, the *nos* promoter was less favored than *zpg* because fecundity was reduced by 45% in crosses between drive heterozygous females and wild type males as well as in crosses between drive heterozygous males and wild type females^18^. We model this as a fitness cost from leaky somatic expression and cleavage of Cas9 in both male and female heterozygotes. However, batch effects may have influenced fecundity determination in this study, because the wild-type controls for each promoter themselves had substantial differences, Further, there is no clear explanation for somatic cleavage negatively affecting male heterozygotes. We therefore also model an alternative interpretation of the nos drive in which there is no somatic fitness cost in males (which we term nosF), which we consider is likely a more accurate representation of this drive. If we further assume that male drive heterozygotes do not have a fitness cost from somatic expression and can serve as a control for female drive heterozygotes, then only a 15% fitness reduction from somatic expression in the females (together with the same 14% embryo resistance allele formation rate in the progeny of female drive heterozygotes) can explain their relative fecundity. We consider this as another alternative possible parameter set (nosF2). We also model a fourth drive in which there are no somatic fitness costs in either sex, which is within the margin of error of the fecundity experiment, though we consider this a highly optimistic parameter set (nosF3).

**Figure S1.**
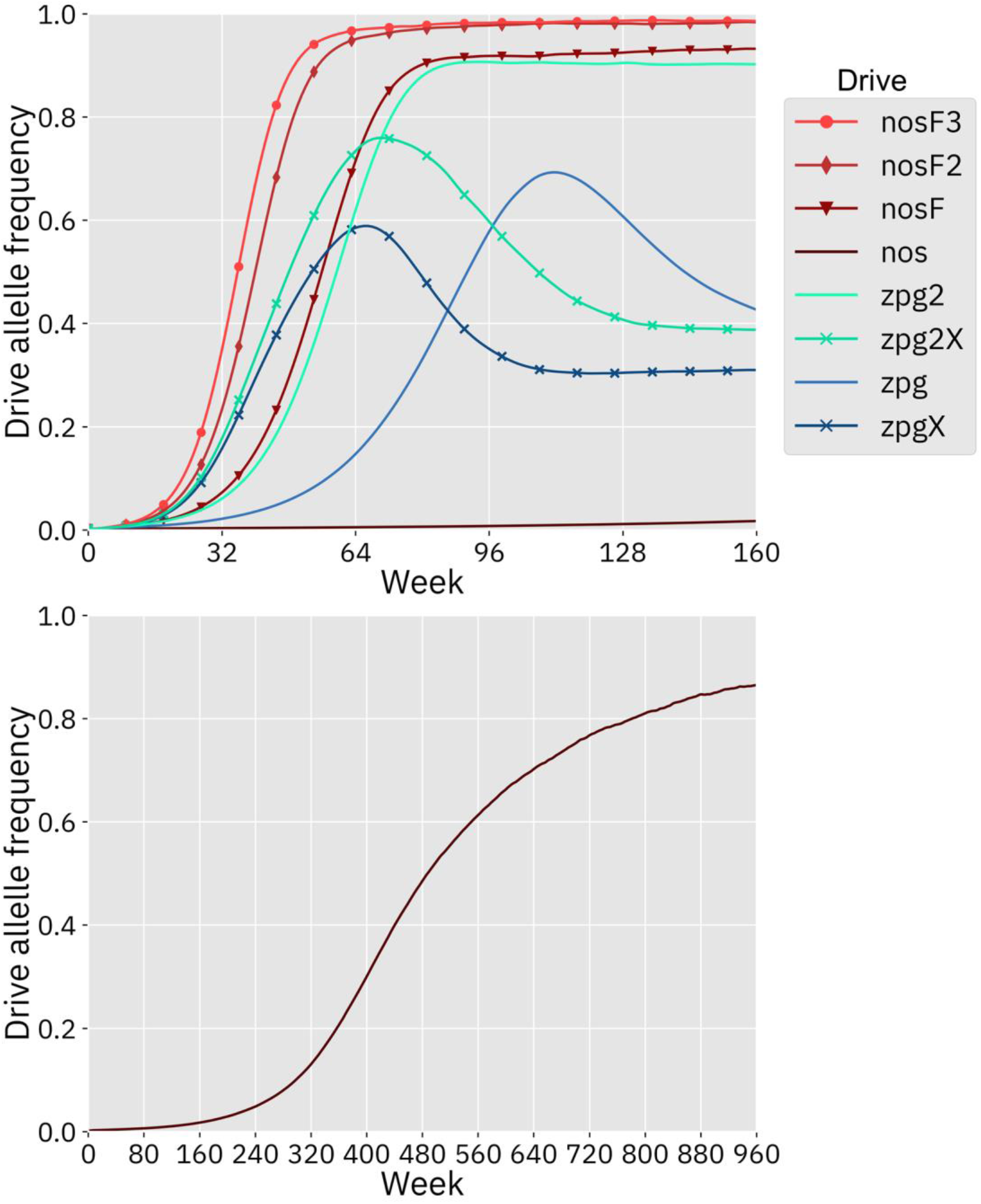
Drive allele frequency trajectories in the mosquito model. Using default parameters and with 20 replicates per drive, each drive was released into a panmictic population. The average allele frequency for each week is displayed. Offspring were artificially generated from fertile individuals at high rates to prevent complete population suppression even at high drive frequencies and genetic loads. The nos drive increases more slowly at low frequency, but eventually reaches a high equilibrium frequency, as seen in the lower panel.

**Figure S2.**
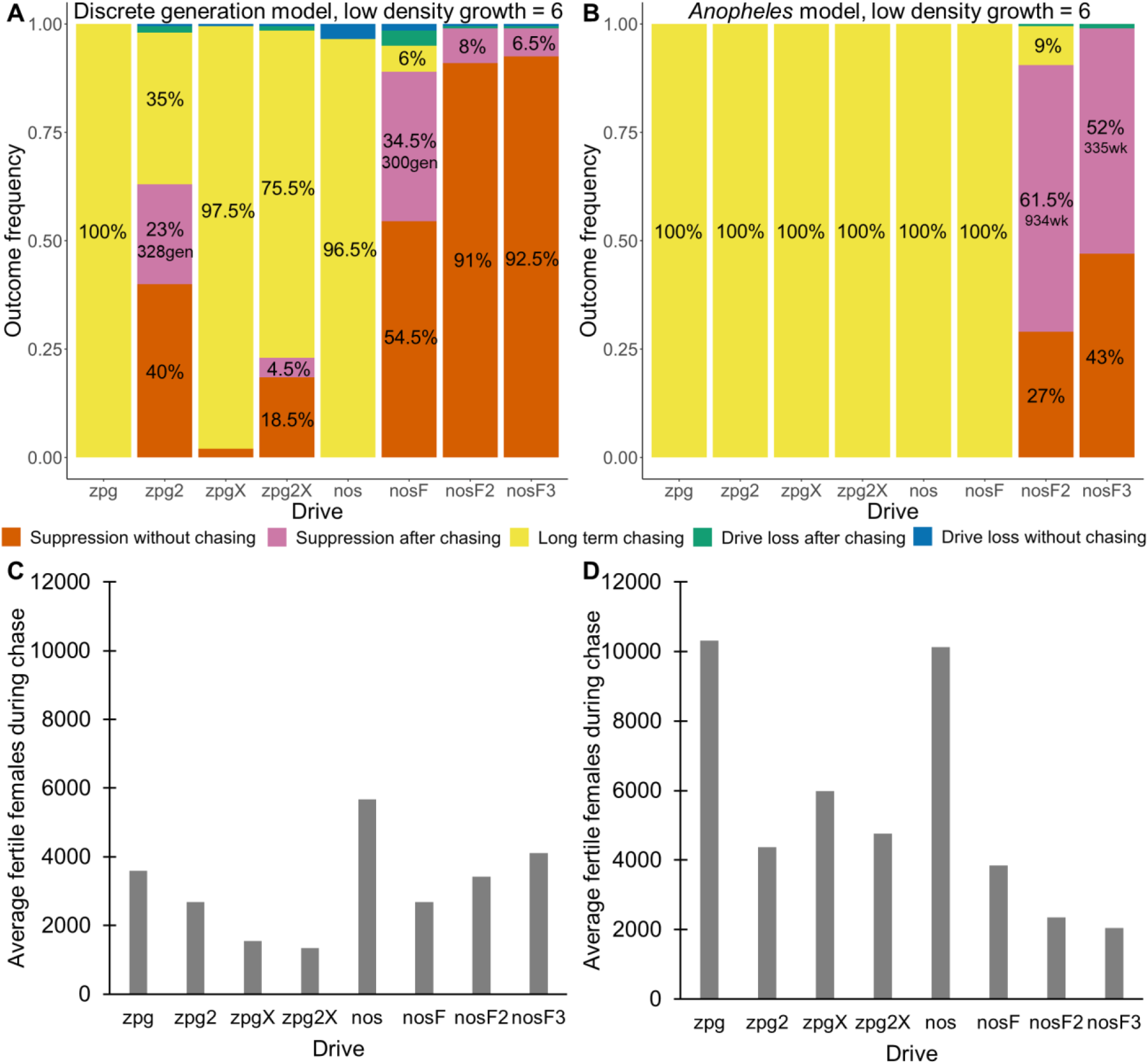
Outcomes in the spatial models with reduced low-density growth rate. Using default parameters, a low-density growth rate of 6, and with 200 replicates per drive, each drive was released into the middle of a wild-type population. The outcome was recorded after 1000 generations or when the population was eliminated for the discrete-generation (**A**) and *Anopheles*-specific (**B**) models. In outcomes involving chasing followed by suppression, the number of generations (gen) or weeks (wk) between the start of chasing and population elimination is shown. Also displayed is the average number of fertile females during periods of chasing (including both long-term and short-term chasing outcomes) for the discrete-generation (**C**) and *Anopheles*-specific (**D**) models. Due to the high number of replicates, the error for each data point is negligible, except for the nosF2 and nosF3 drives in the discrete-generation model due to the short duration of chasing.

**Figure S3.**
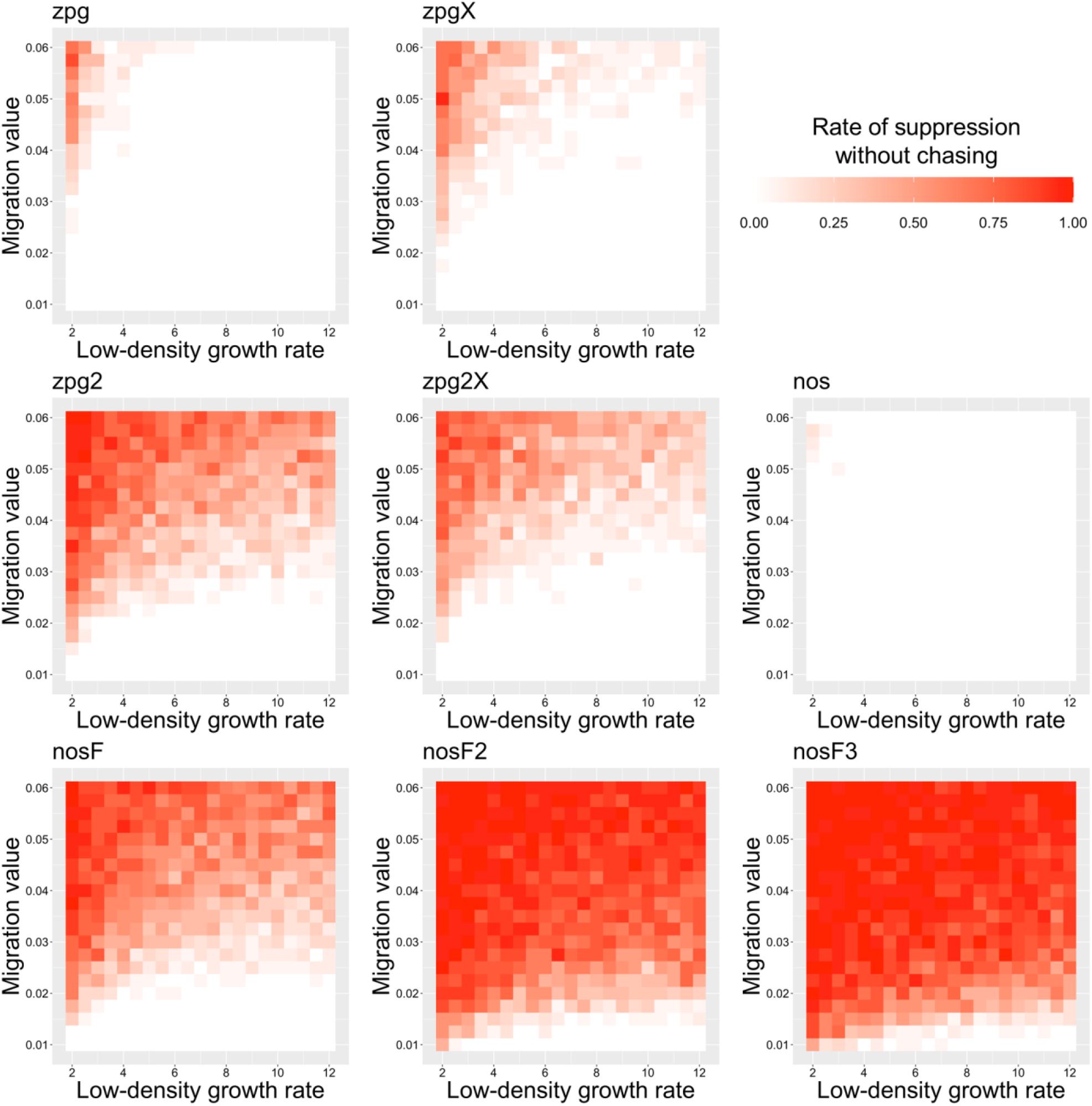
The rate of suppression without chasing in the discrete-generation model. Drive-carrying individuals were released into the middle of a wild-type population in a 1×1 area. The proportion of simulations in which suppression occurred either before a chase or within 10 generations of the start of chasing is shown. Each point represents the average of 20 simulations.

**Figure S4.**
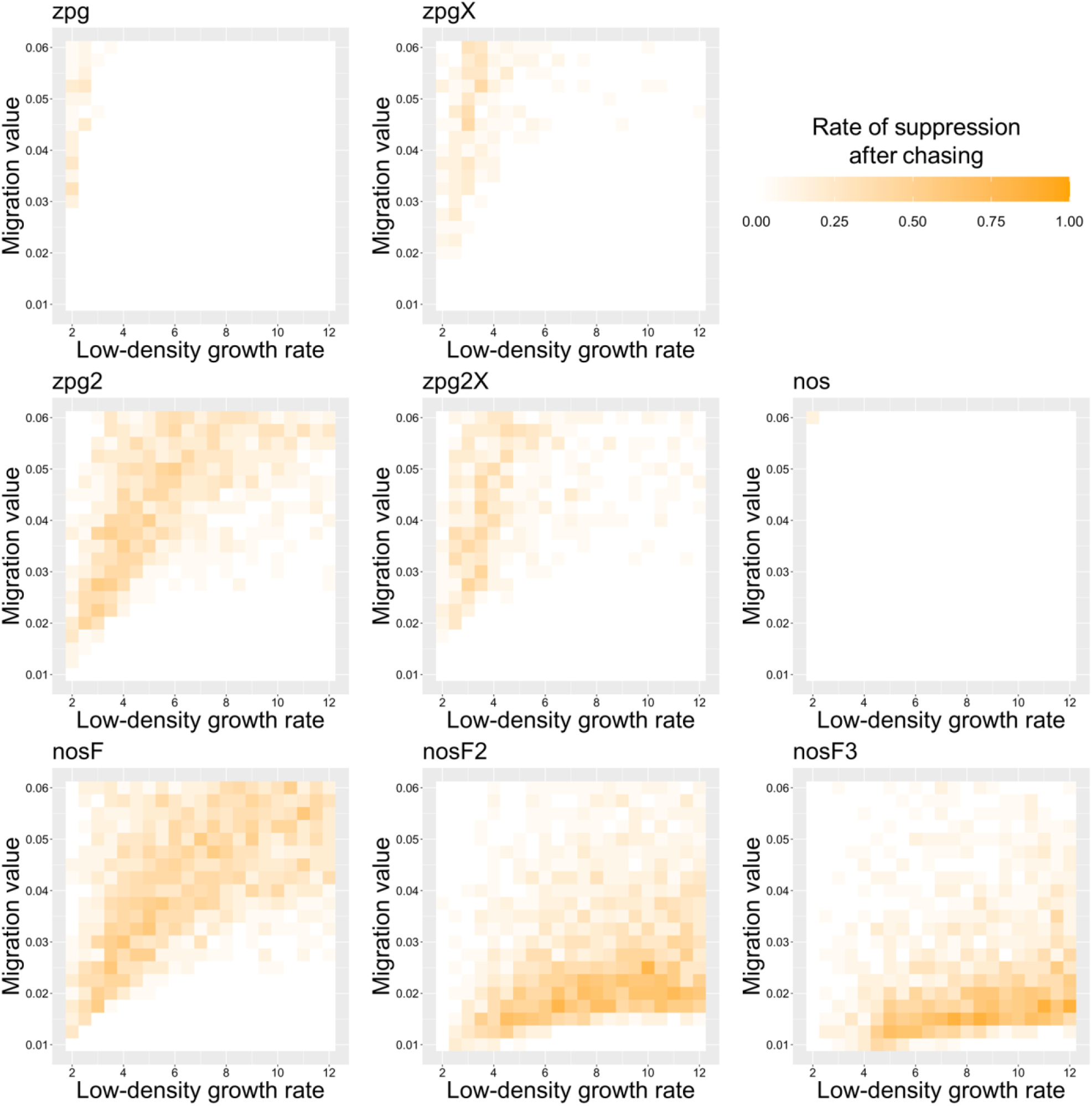
The rate of suppression after chasing in the discrete-generation model. Drive-carrying individuals were released into the middle of a wild-type population in a 1×1 area. The proportion of simulations in which suppression occurred after a chase that lasted a minimum of 10 generations is shown. Each point represents the average of 20 simulations.

**Figure S5.**
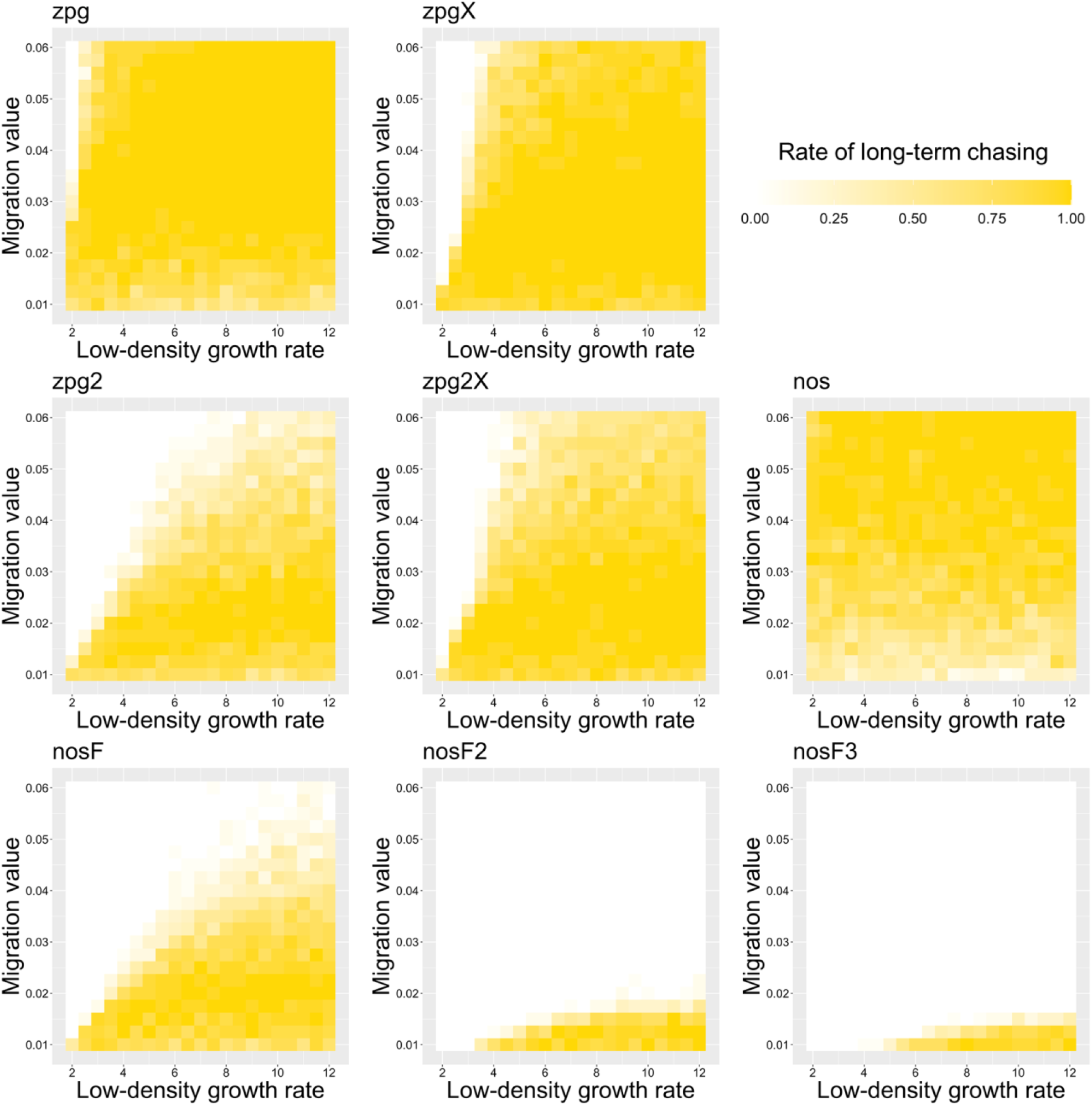
The rate of long-term chasing in the discrete-generation model. Drive-carrying individuals were released into the middle of a wild-type population in a 1×1 area. The proportion of simulations in which a long-term chasing outcome (defined by a chase continuing for 1000 generations after drive release) occurred is shown. Each point represents the average of 20 simulations.

**Figure S6.**
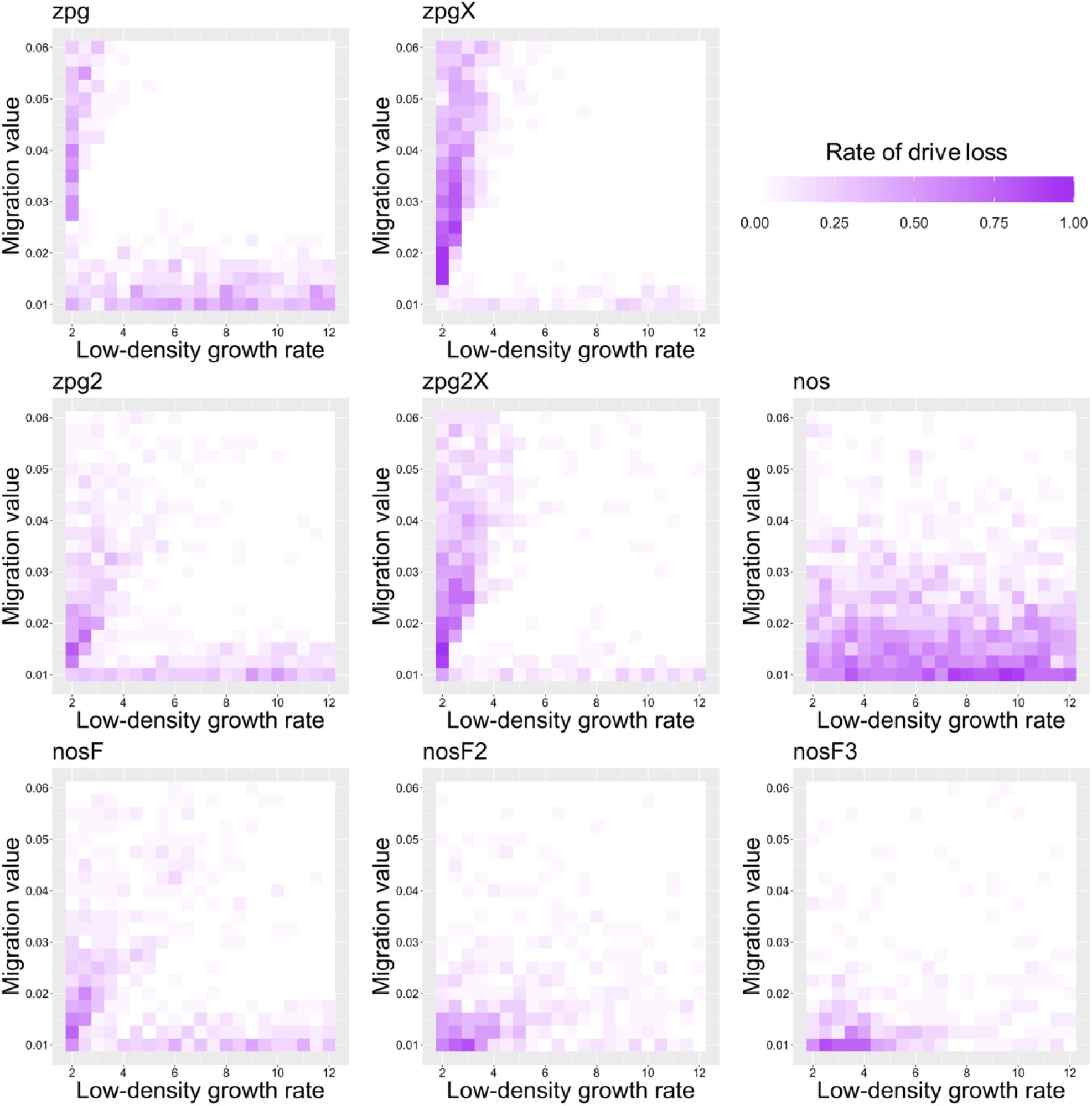
The rate of drive loss in the discrete-generation model. Drive-carrying individuals were released into the middle of a wild-type population in a 1×1 area. The proportion of simulations in which the drive was lost from the population is shown. Each point represents the average of 20 simulations.

**Figure S7.**
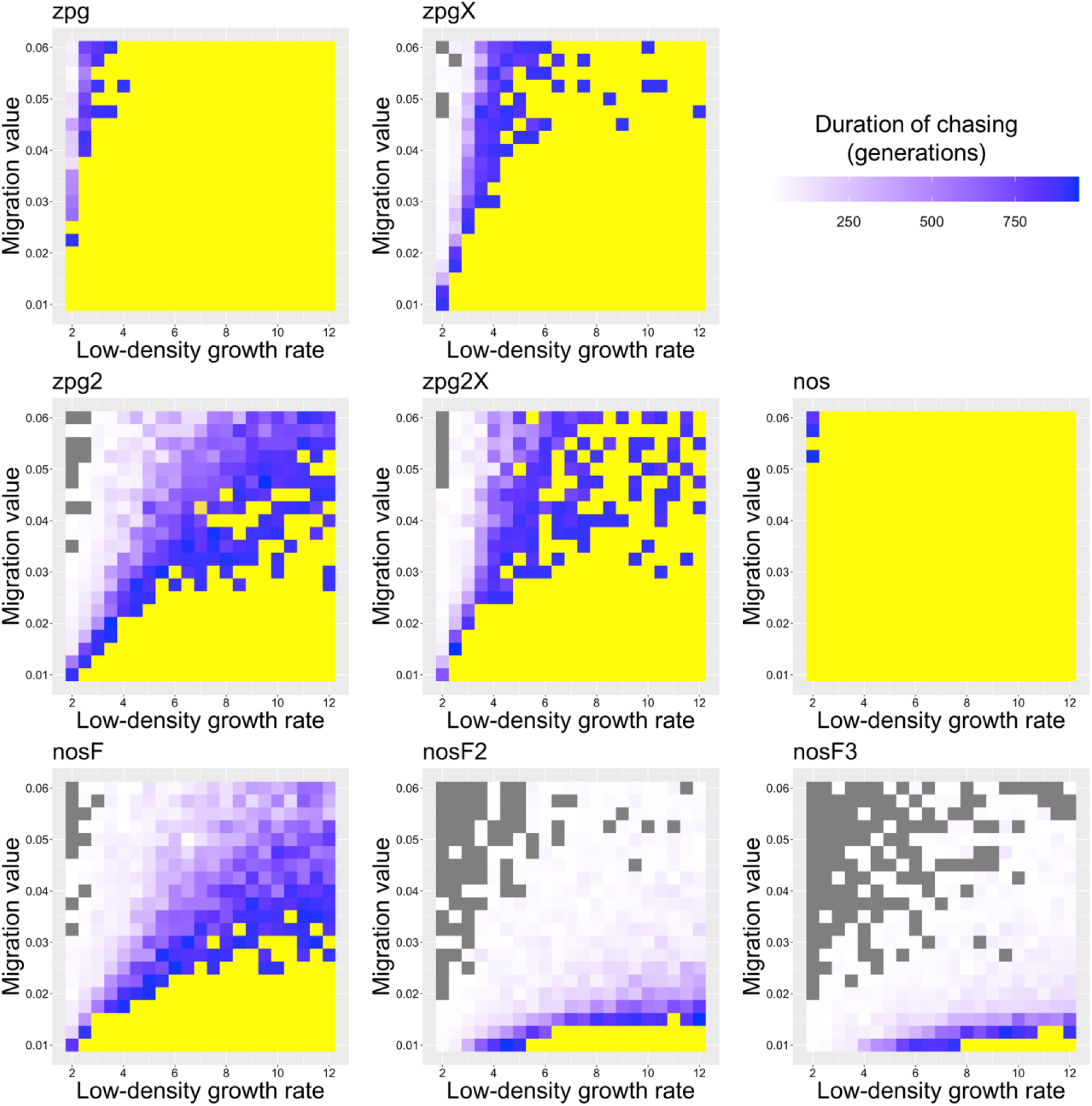
The duration of chasing prior to suppression in the discrete-generation model. Drive-carrying individuals were released into the middle of a wild-type population in a 1×1 area. The number of generations between the start of chasing and population elimination is shown. Each point represents the average of 20 simulations. Grey represents parameter combinations in which chasing did not occur in any simulation, and yellow represents parameter combinations in which chasing occurred but did not end in suppression in any simulation.

**Figure S8.**
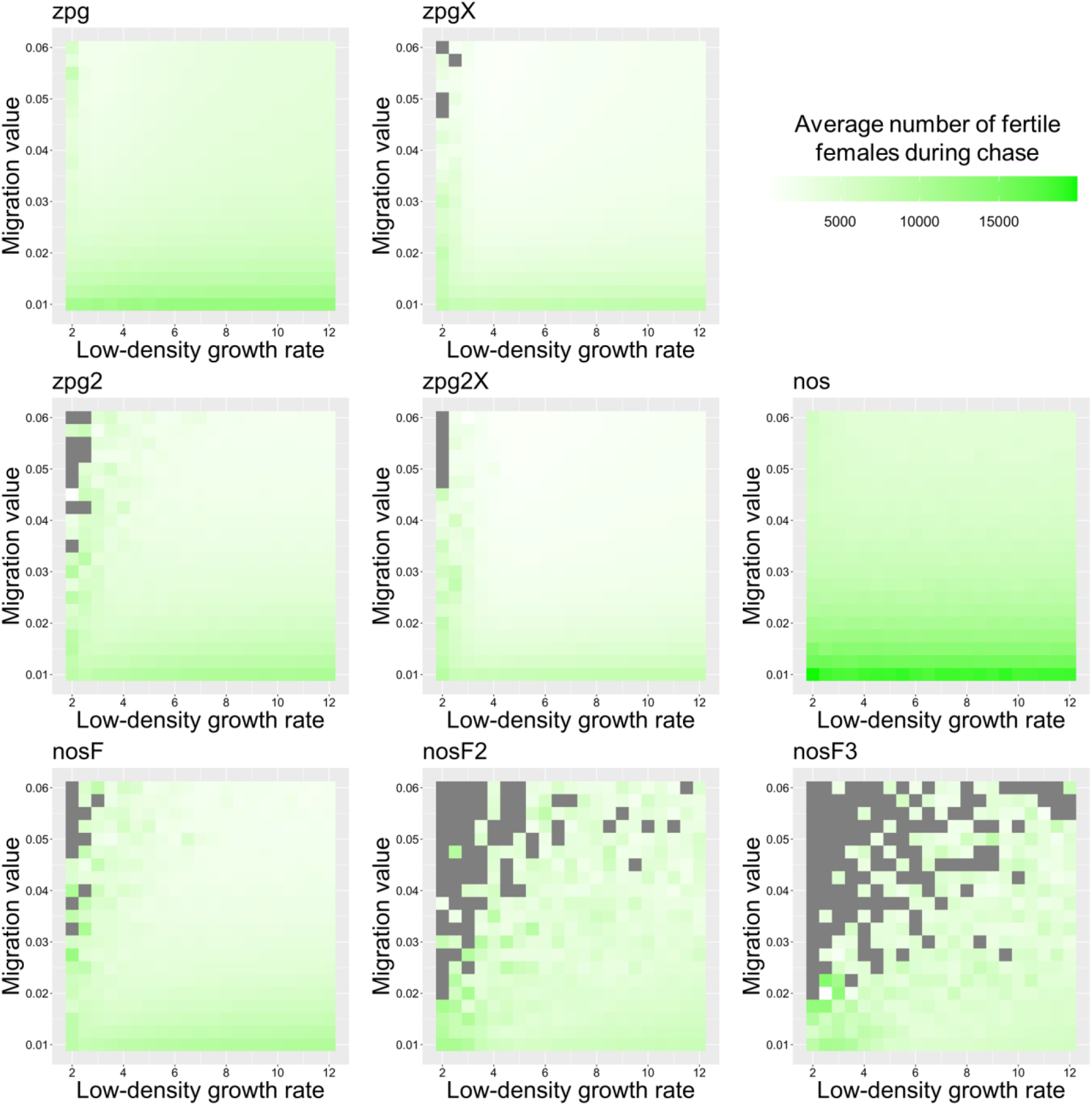
The average number of fertile females during chasing in the discrete-generation model. Drive-carrying individuals were released into the middle of a wild-type population in a 1×1 area inhabited by an average of 25,000 females. The average number of fertile females during periods of chasing is shown. Each point represents the average of 20 simulations. Grey represents parameter combinations in which chasing did not occur in any simulation.

**Figure S9.**
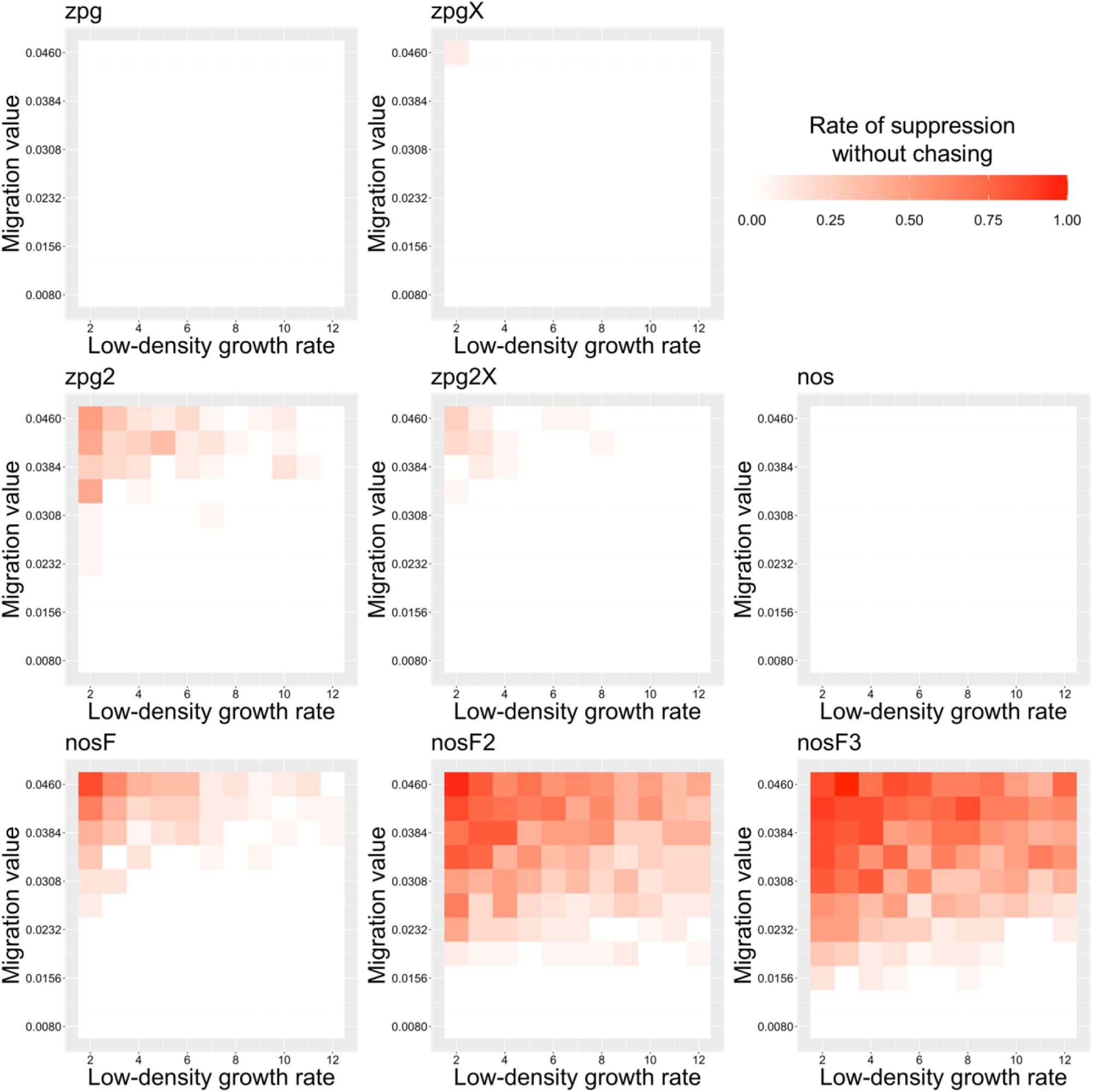
The rate of suppression without chasing in the *Anopheles* model. Drive-carrying mosquitoes were released into the middle of a wild-type population in a 1×1 area. The proportion of simulations in which suppression occurred either before a chase or within 32 weeks of the start of chasing is shown. Each point represents the average of 20 simulations.

**Figure S10.**
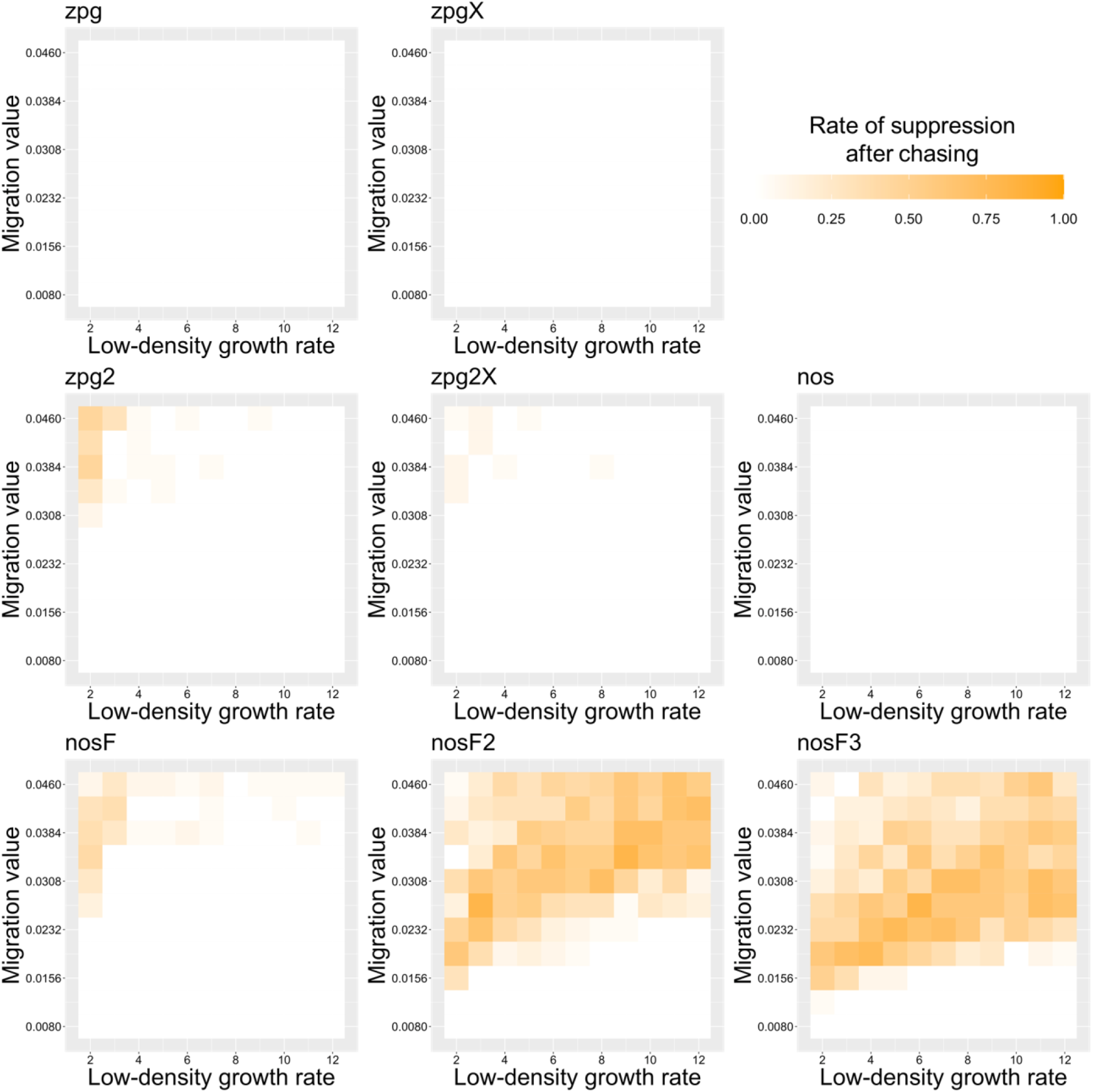
The rate of suppression after chasing in the *Anopheles* model. Drive-carrying mosquitoes were released into the middle of a wild-type population in a 1×1 area. The proportion of simulations in which suppression occurred after a chase that lasted a minimum of 32 weeks is shown. Each point represents the average of 20 simulations.

**Figure S11.**
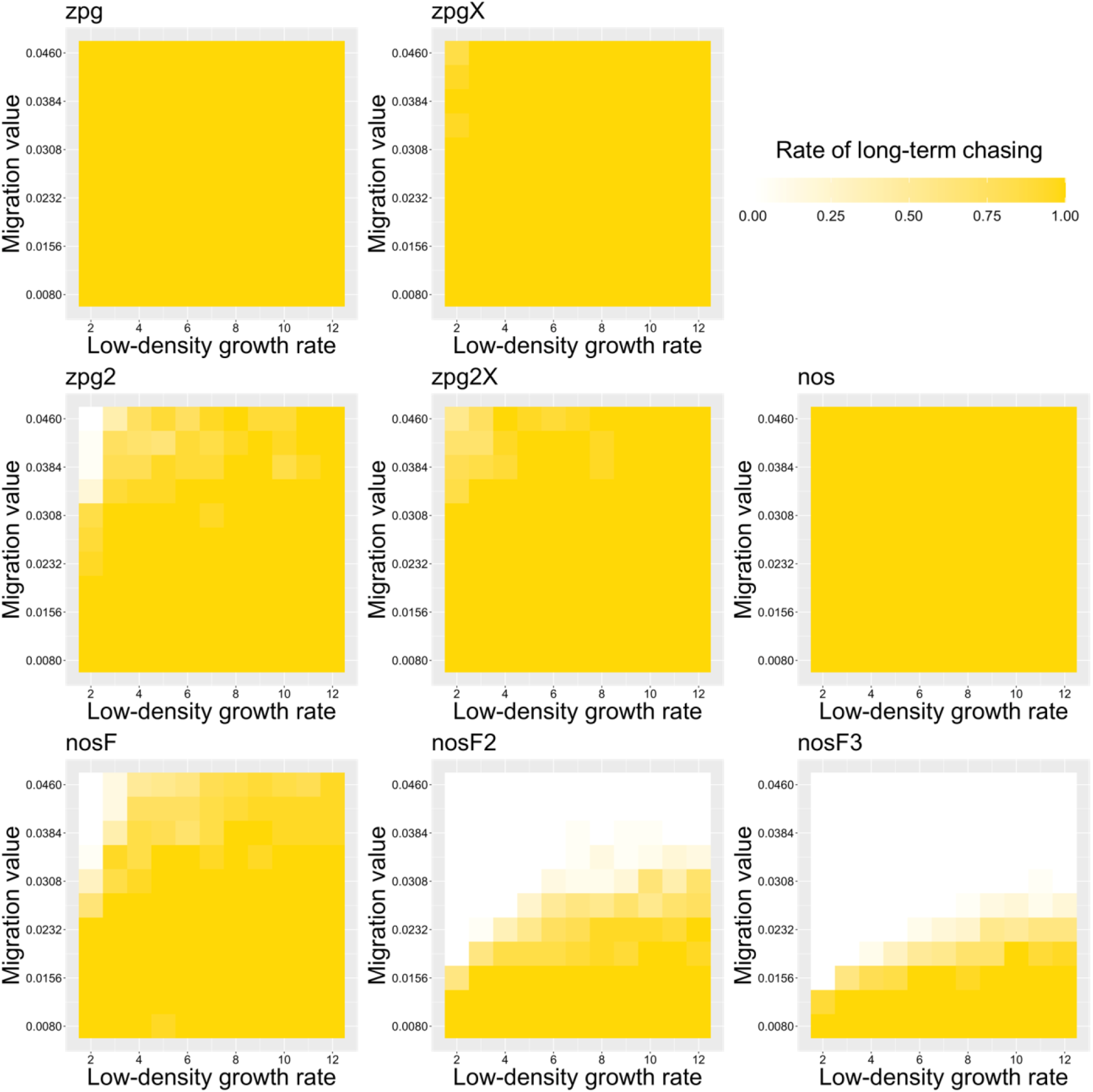
The rate of long-term chasing in the *Anopheles* model. Drive-carrying mosquitoes were released into the middle of a wild-type population in a 1×1 area. The proportion of simulations in which a long-term chasing outcome (defined by a chase continuing for 3167 weeks after drive release) occurred is shown. Each point represents the average of 20 simulations.

**Figure S12.**
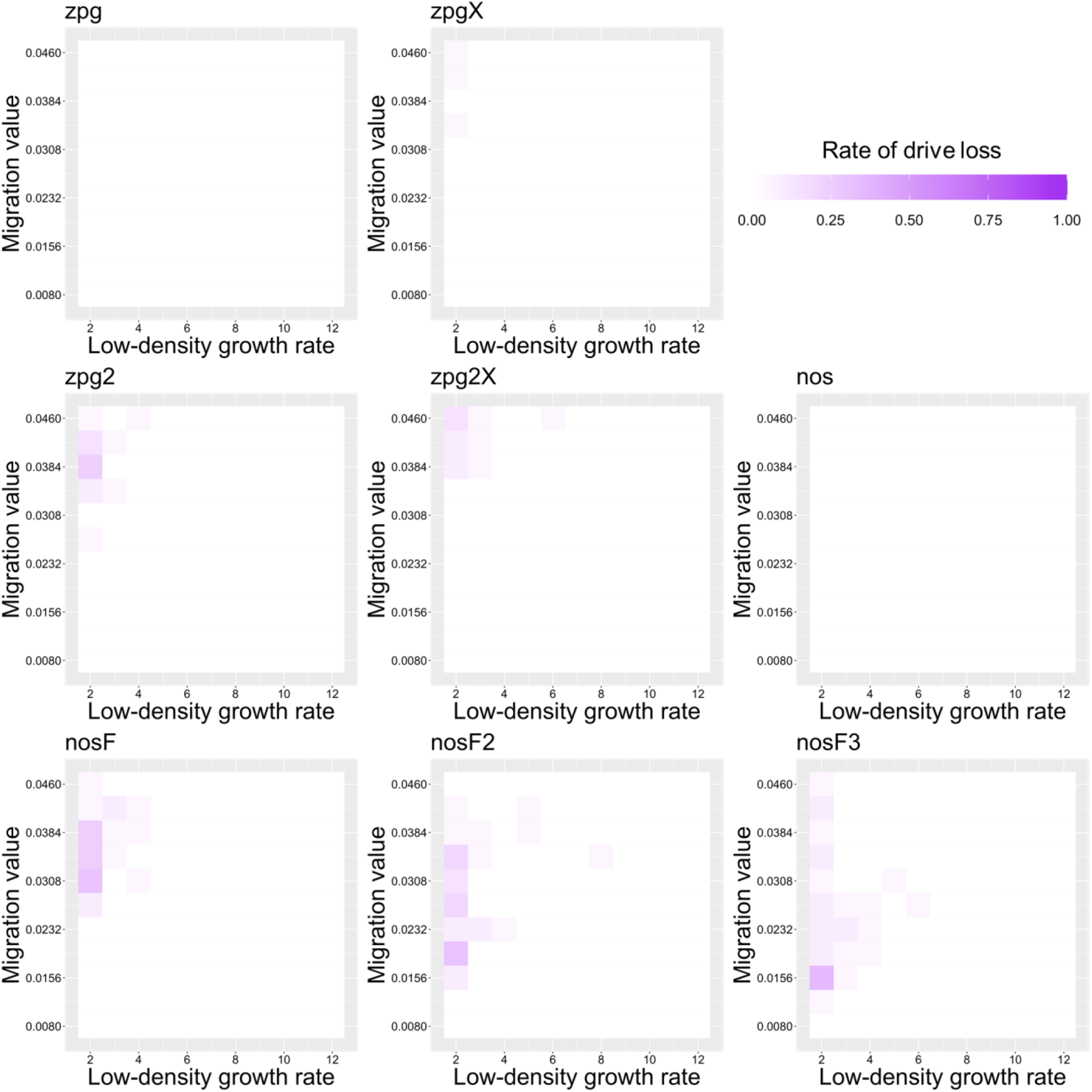
The rate of drive loss in the *Anopheles* model. Drive-carrying mosquitoes were released into the middle of a wild-type population in a 1×1 area. The proportion of simulations in which the drive was lost from the population is shown. Each point represents the average of 20 simulations.

**Figure S13.**
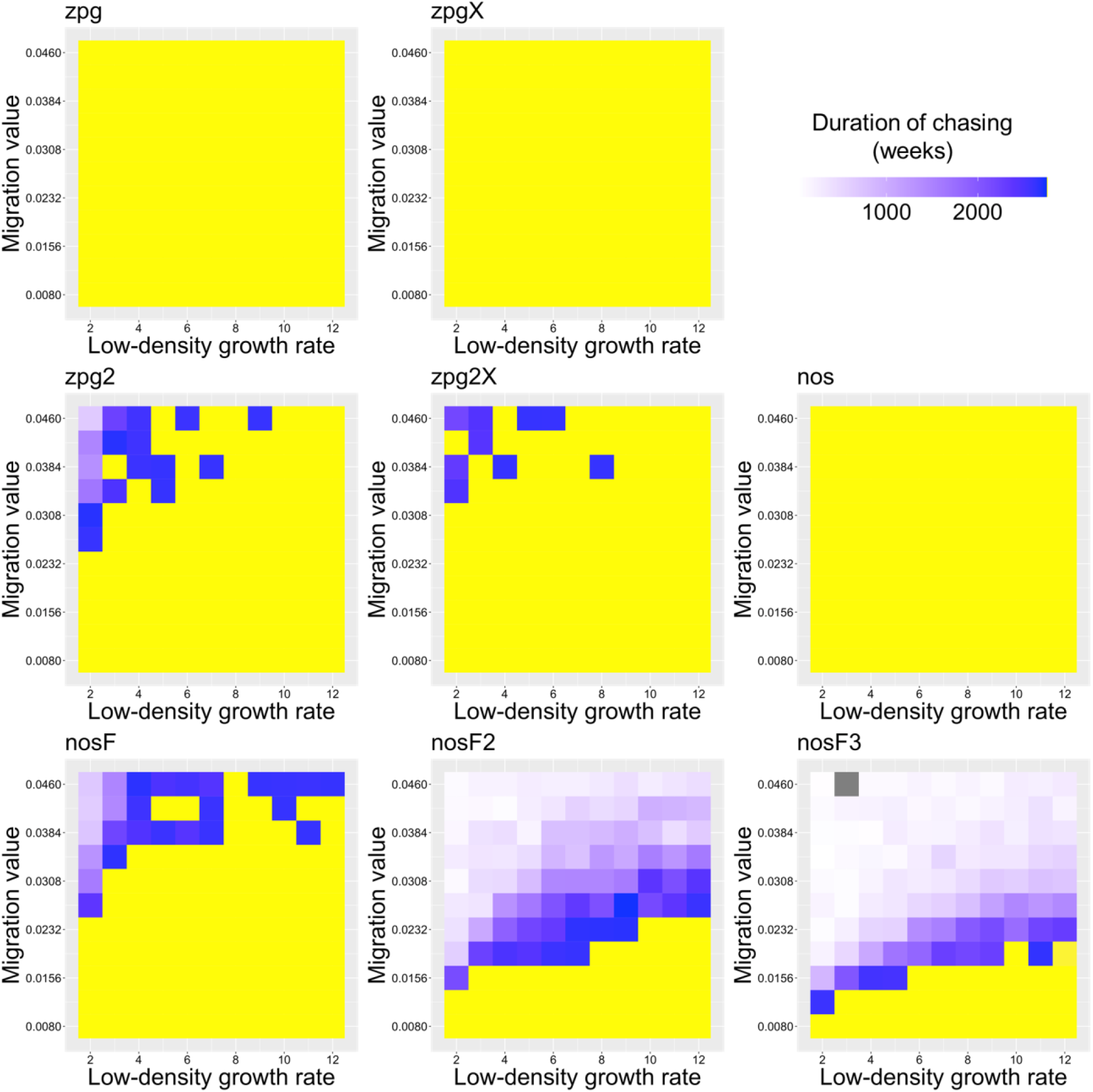
The duration of chasing prior to suppression in the *Anopheles* model. Drive-carrying mosquitoes were released into the middle of a wild-type population in a 1×1 area. The number of generations between the start of chasing and population elimination is shown. Each point represents the average of 20 simulations. Grey represents parameter combinations in which chasing did not occur in any simulation, and yellow represents parameter combinations in which chasing occurred but did not end in suppression in any simulation.

**Figure S14.**
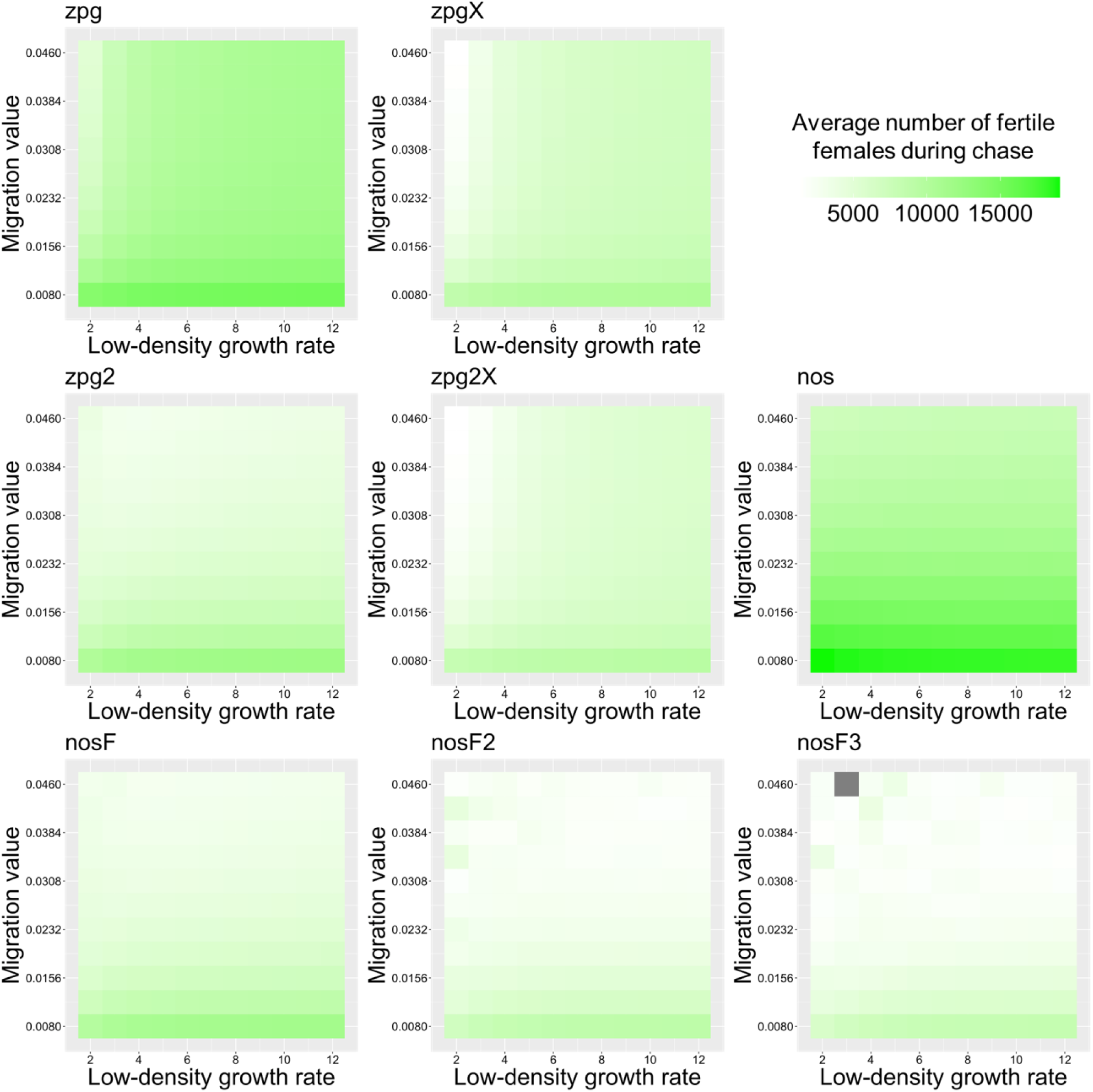
The average number of fertile females during chasing in the *Anopheles* model. Drive-carrying mosquitoes were released into the middle of a wild-type population in a 1×1 area inhabited by an average of 25,000 females. The average number of fertile females during periods of chasing is shown. Each point represents the average of 20 simulations. Grey represents parameter combinations in which chasing did not occur in any simulation.

